# The genomes and epigenomes of aquatic plants (Lemnaceae) promote triploid hybridization and clonal reproduction

**DOI:** 10.1101/2023.08.02.551673

**Authors:** Evan Ernst, Bradley Abramson, Kenneth Acosta, Phuong T.N. Hoang, Cristian Mateo-Elizalde, Veit Schubert, Buntora Pasaribu, Nolan Hartwick, Kelly Colt, Anthony Aylward, Seung Cho Lee, Umamaheswari Ramu, James A. Birchler, Ingo Schubert, Eric Lam, Todd P. Michael, Robert A. Martienssen

## Abstract

The Lemnaceae (duckweeds) are the world’s smallest but fastest growing flowering plants. Prolific clonal propagation facilitates continuous micro-cropping for plant-based protein and starch production, and holds tremendous promise for sequestration of atmospheric CO_2_. Here, we present chromosomal assemblies, annotations, and phylogenomic analysis of *Lemna* genomes that uncover candidate genes responsible for the metabolic and developmental traits of the family, such as anatomical reduction, adaxial stomata, lack of stomatal closure, and carbon sequestration via crystalline calcium oxalate. Lemnaceae have selectively lost genes required for RNA interference, including Argonaute genes required for reproductive isolation (the triploid block) and haploid gamete formation. Triploid hybrids arise commonly among *Lemna*, and we have found mutations in highly-conserved meiotic crossover genes that could support polyploid meiosis. Syntenic comparisons with *Wolffia* and *Spirodela* reveal that diversification of these genera coincided with the “Azolla event” in the mid-Eocene, during which aquatic macrophytes reduced high atmospheric CO_2_ levels to those of the current ice age.

Facile regeneration of transgenic fronds from tissue culture, aided by reduced epigenetic silencing, makes *Lemna* a powerful biotechnological platform, as exemplified by recent engineering of high-oil *Lemna* that outperforms oil seed crops.

## Introduction

The Lemnaceae ^1^ are a family of freshwater aquatic macrophytes commonly known as duckweeds ^2^ and are sometimes referred to as water lentils and watermeal. The Lemnaceae reproduce by reiterative vegetative budding from a “pocket” of meristematic stem cells, doubling once per day under optimal conditions. Free floating clonal reproduction provides the optimal environment for rapid plant growth, and the Lemnaceae have the shortest biomass doubling time of any flowering plant, making them attractive for micro-farming, and for nitrate, phosphate, and CO_2_ remediation. However, they are true flowering plants, and some species in each of the five genera (*Spirodela, Landoltia, Lemna, Wolffia,* and *Wolffiella*) can produce simple flowers and fruits with 1-5 seeds in response to hormones, nutrients, temperature and daylength (Figure 1A). The clonal growth habit allows a high frequency of polyploidy as well as interspecific hybridization ^3–5^, suggesting that reproductive isolation barriers in the seed were lost in the absence of obligate sexual reproduction ^3^. In particular, *L. turionifera* (T) and *L. minor* (M) form frequent polyploid hybrids, known as *L. japonica* ^5^, which have enhanced vigor compared with diploid relatives under certain environmental conditions ^6^, perhaps explaining their adventitious selection for biotechnological applications ^7,8^. Isolates classified as either *L. minor* or *L. japonica* are difficult to distinguish morphologically or by plastid markers ^9^, though genetic barcoding with polymorphic nuclear markers confirms their classification as distinct species ^4^.

**Figure 1.**
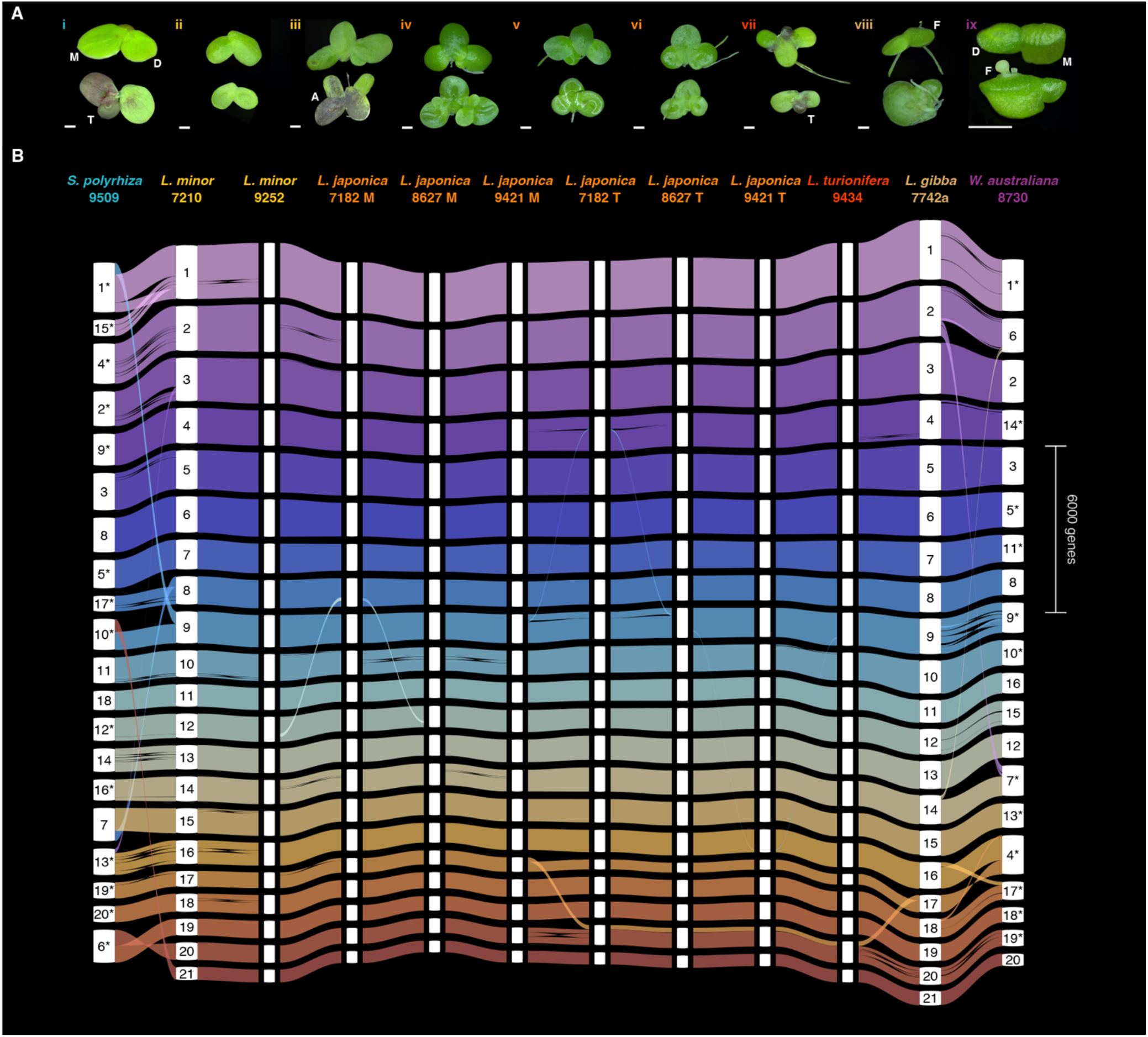
Lemnaceae habit and genomes. **(A)** Species in this study. i: *Spirodela polyrhiza* 9509; ii: *Lemna minor* 7210; iii: *Lemna minor* 9252; iv: *Lemna japonica* 7182; v: *Lemna japonica* 8627; vi: *Lemna japonica* 9421; vii: *Lemna turionifera* 9434; viii: *Lemna gibba* 7742a; ix:*Wolffia australiana* 8730. Darkfield microscopy of colonies of mother fronds (M) bearing clonal daughter frond progeny (D) (1mm scale bar). Turions (T) are visible after 40 days of growth in dilute media in *L. turionifera* (vii) colonies, but not in those of *L. minor* (ii-iii) or *L. japonica* (iv-vi). *S. polyrhiza* (i) produced turions on dilute media. Starvation elicited a strong anthocyanin (A) response in turion-producing plants and in the fronds of *L. minor* 9252 (iii). Flowers (F) are visible in *L. gibba* (viii) and *W. australiana* (ix) after growth on inductive media. **(B)** Gene-level synteny. The genomes and subgenomes of *Lemna* and *Wolffia* species were sequenced using long-read single molecule sequencing, assembled into 21 (*Lemna*) or 20 (*Wolffia*) pseudomolecules with chromatin conformation capture, and annotated for direct comparison of gene content. *Lemna* chromosomes were numbered by size in *L. minor* 7210 (common duckweed). Ribbons represent blocks of syntenic protein-coding gene loci. * = chromosomes inverted relative to their reference representation to more clearly show syntenic relationships.

Here, we use single molecule nanopore sequencing and Hi-C contact mapping to generate chromosome-resolved genome assemblies of *Lemna* species, including the first assemblies of *L. japonica* interspecific hybrids, revealing that they form with variable parental dosage as diploids and reciprocal triploids. Genome in situ hybridization (GISH) using super-resolution microscopy confirms these chromosomal assignments. Comparisons between *Lemna, Wolffia* and *Spirodela* allow syntenic relationships to be determined across the Lemnaceae, and have enabled us to consistently number and orient the *Lemna* chromosome sets. Chromatin architecture, tandem repeat distribution, and chromatin interaction mapping indicates unexpected association between the sub genomes, which could be indicative of holocentricity. We also determine patterns of DNA methylation as well as small RNA accumulation. We find that the loss of small RNA and genes required for RNA dependent DNA methylation could account for the high frequency of polyploids in Lemnaceae. This is because RNA dependent DNA methylation is required in flowering plants for the “triploid block”: a reproductive barrier in which triploid seeds abort ^10,11^.

Analysis of gene content reveals highly diverged and missing orthogroups, which include candidate genes for reduced morphology, altered metabolic profiles, and signaling pathways. Master transcription factors as well as key enzymes missing in the Lemnaceae may account for the loss of lateral roots and root hairs, altered lipid and lignin accumulation, and the loss of stomatal response to elevated CO_2_. Floating freshwater aquatic ferns related to *Azolla* may have been responsible for the most dramatic historical reduction in atmospheric CO_2_ levels in the late Eocene ^12^, and would have greatly benefited from the ability to maintain open stomatal apertures under elevated CO_2_ concentrations. Efficient regeneration of transgenic fronds from tissue culture, and reduced epigenetic silencing, could make *Lemna* a powerful biotechnological platform, and the Lemnaceae offer a unique opportunity to engineer CO_2_ capture and sequestration, as well as biofuel production, in the modern age. A recent study showed *L. japonica* plants simultaneously overexpressing *WRINKLED1*, *DIACYLGLYCEROL ACYLTRANSFERASE*, and *OLEOSIN* can accumulate oil at up to 8.7% of dry mass, illustrating the potential for metabolic pathway manipulation in Lemnaceae ^13^.

## Results

### Chromosome-resolved Lemnaceae genome assemblies

#### Genome architectures and synteny

We used Oxford Nanopore Technologies (ONT) single molecule long reads, paired with high-throughput chromatin conformation capture (Hi-C) contact mapping or reference-based scaffolding, to generate chromosome-resolved *de novo* genome assemblies for 8 duckweed accessions representing 5 species, and protein coding gene annotations spanning 3 genera (Figure 1, Table S1, Figure S1). Mean raw ONT read coverage ranged from 38x to 105x (Table S2), and contig N50s varied from 3.2 Mbp - 13.9 Mbp (Table S1). *Lemna japonica* 8627 (previously classified as *Lemna minor*) was among our initial targets for whole genome sequencing due to its amenability to genetic transformation and use as a recombinant expression platform ^14,15^. Individual ONT reads are long enough to span distant tracts of single nucleotide and structural variants (SNVs and SVs), enabling the separation of two homeologous chromosome sets in draft assemblies of this accession. This prompted us to sequence three additional *L. japonica* accessions and their founder species to better understand hybridization and genome variation within the genus (Figure 2A).

**Figure 2.**
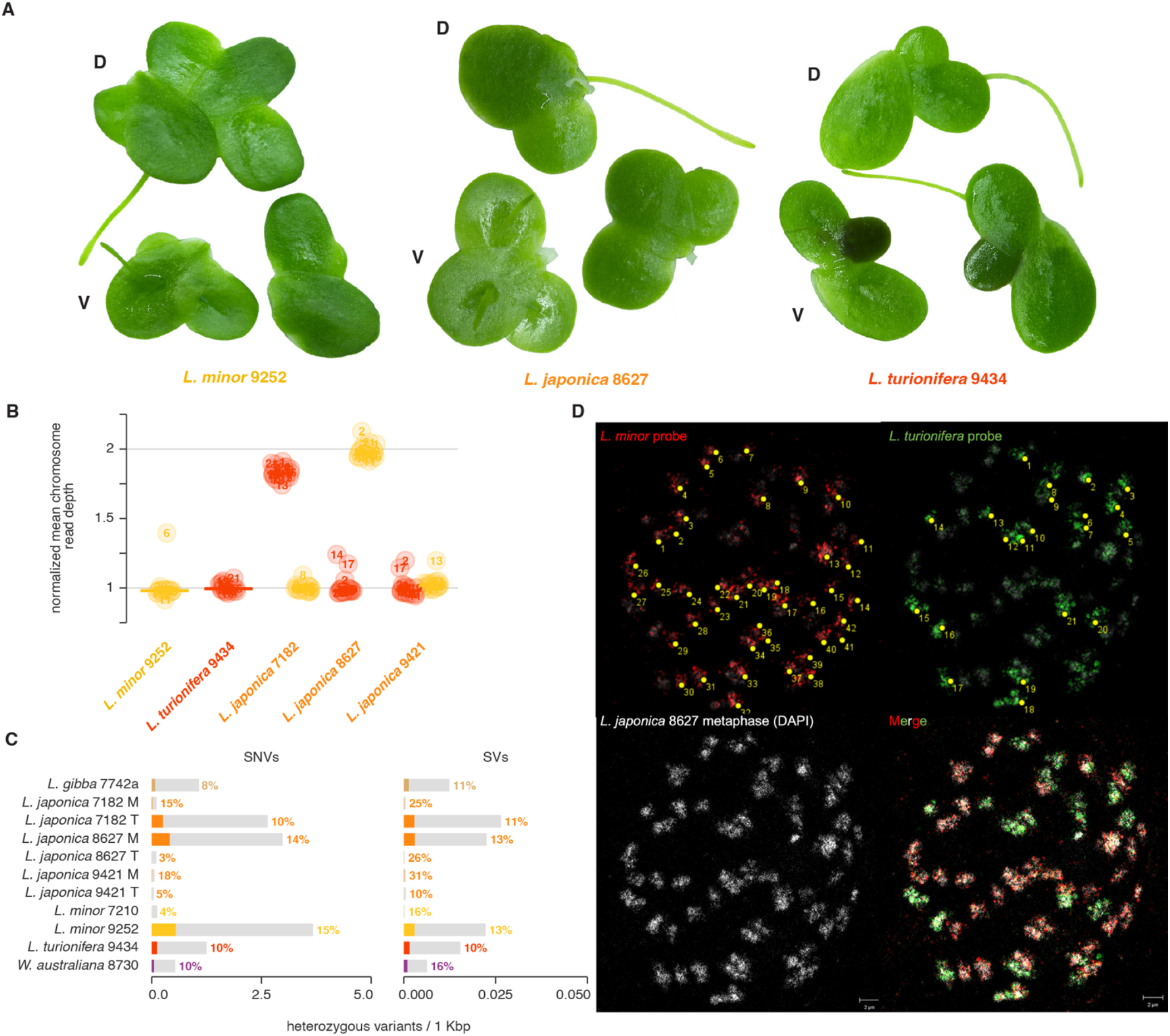
Subgenome dosage and heterozygosity in diploid and triploid L. japonica. **(A)** Dorsal (D) and ventral (V) views of typical *L. minor, L. japonica*, and *L. turionifera* frond colonies. Turions are forming in one of the two meristematic recesses of *L. turionifera*. **(B)** Read depth per pseudomolecule in *L. japonica* hybrids and representative diploid parent assemblies, normalized to the mode of read depth across each subgenome assembly. Pseudomolecules are numbered as in Figure 1. **(C)** Heterozygous variant calls made from long reads mapped back to each haplotype-collapsed assembly. Single nucleotide variants and INDELs < 50 bp (SNVs), and structural variants > 50 bp (SV) counts per kilobase of reference genome sequence are shown. The subgenomes of *L. japonica* hybrids are separately indicated by their *L. minor* (M) or *L. turionifera* (T) karyotypes. Variants overlapping protein coding gene annotations are highlighted with color and their proportion is shown to the right of the bars. SNV and SV levels in diploid subgenomes reflect substantial heterozygosity, except for Lm7210, where levels are at the false positive rate observed in monoploid subgenomes 7182 M, 8627 T, 9421 M, and 9421 T. Lm7210 thus appears to be a doubled haploid. **(D)** Genome in-situ hybridization (GISH) of genomic DNA of *L. minor* (in red) and *L. turionifera* (in green) on metaphase chromosomes of *L. japonica* 8627 visualized by 3D-structured illumination microscopy.

We resolved M (*L. minor*) and T (*L. turionifera*) subgenomes into two haplotype-collapsed sets of 21 chromosomes for Lj7182, Lj8627, and Lj9421, confirming that the *L. japonica* taxon represents distinct interspecific hybrids of *L. minor* and *L. turionifera* (Figures 1A, 1B, and 2B). Genomic read mapping was used to assess dosage of each parental haplotype between the three accessions, and together with nuclear genome size estimates by flow cytometry this indicated that hybrids form both as MT diploid (Lj9421) and reciprocal MTT (Lj7182) and MMT (Lj8627) triploids (Figures 2B and S2). Whole genome alignment and synteny mapping supported a consistent *L. turionifera* karyotype distinguished from *L. minor* and the more distantly related *L. gibba* by the translocation of 3.5 Mbp of one arm of Chr17 to Chr20 (Figure 1B). Further highlighting the difficulty in discriminating *L. minor* from *L. japonica* hybrids, the assembly of accession 9252, originally labeled *L. japonica*, lacked an *L. turionifera* subgenome, consistent with a previous report that it is a heterozygous diploid *L. minor* ^4^. In contrast, Lm7210 from South Africa, had a rate of heterozygous short variant calls comparable to the single-copy subgenomes of the hybrids, which represents the false discovery rate indicating that Lm7210 could be a natural doubled haploid (Figure 2C). Although heterozygosity was evident in the other diploid genomes (Figure 2c), it was very low compared with terrestrial plants, as it is in *Spirodela*, and other aquatic plants ^16–19^. The genome dosage and parental composition of Lj8627 was independently confirmed by Genome *in situ* Hybridization (GISH) using the *L. minor* and *L. turionifera* genomes as probes (Figure 2D). Structured illumination microscopy (SIM) was used to resolve individual metaphase chromosome pairs from each subgenome, revealing 21 chromosomes from *L. turionifera* and 42 chromosomes from *L. minor*. We note that metaphase chromosomes from each subgenome had no obvious centromeric constriction.

#### Ribosomal DNA (rDNA) repeats are rearranged in triploid hybrids

In most cases we were able to determine the chromosomal locations of highly-conserved rDNA repeat arrays which showed evidence of karyotypic plasticity (Figure S3). DNA FISH studies detected just one 45S rDNA locus in the majority of duckweed accessions surveyed, yet some *Wolffia* and *Wolffiella* clones had two loci ^20^. In the case of *W. australiana* 8730 (Wa8730), a sole intact locus was assembled on Chr14, which was homologous to *Lemna* Chr4. A remnant locus was also present on Wa8730 Chr4, which was homeologous to Chr16, Chr17 and Chr20 in *Lemna spp.*, near the Chr17:Chr20 fusion breakpoint (Figures 1B, S1B). In the diploid *L. minor* accessions assembled here, the intact array was located at the end of Chr20, while in the *L. turionifera* subgenomes of the hybrids, the best-conserved array was translocated to Chr17. This was consistent with an rDNA array on the ancestral homeolog of Wa8730 Chr4 migrating to Chr20 in *L. minor*, and Chr17 in *L. turionifera* lineages. The translocation of the Chr17 terminus to Chr20 was shared in all *L. turionifera* assemblies suggesting this may have occurred at the same time (Figures 1B, S1B). However, among the *L. japonica* hybrids, intact and likely active rDNA repeats were assembled at distinct positions. In the MTT hybrid 7182, a conserved array was assembled only at Chr17T, and the remaining loci, including Chr20M, are degraded. In the MMT hybrid 8627, only remnant arrays were present on Chr17T and three other locations but the highly-conserved array sequence was unanchored, as it was in *L. turionifera* Lt9434. Only in the case of the diploid 9421, intact rDNA copies were assembled on both subgenomes. In addition, a consistently degraded remnant locus appeared on Chr6 in the diploids Lm9252 and Lt9434, and on both M and T subgenomes of the *L. japonica* hybrids. Thus active rDNA array degradation occurred in triploid but not diploid hybrids, possibly reflecting dosage of the parental chromosomes on which they reside. Similar rearrangements are frequent in other examples of polyploid hybrids ^21^.

#### Centromere identification and characterization

In most plant genomes the epigenetically defined centromere region is nested in large high copy number tandem duplication (HCNTD) repeat arrays with a typical base repeat length between 150 and 250 base pairs (bp) that form higher order repeats (HOR) ^22,23^. In the model dicot *Arabidopsis thaliana*, these arrays are megabases in length and highly conserved, making them difficult to sequence ^24^. However, while candidate HCNTD were identified in *S. polyrhiza* ^22^, the chromosome resolved genome did not contain this HCNTD ^19,25^. The same is true in the *W. australiana* genome; a HCNTD is present, but these are not large arrays in the genome ^26^. We searched the *Lemna* genomes for HCNTDs, and found that *L. gibba* also lacks a prominent array (Figure 3A). In contrast, *L. minor* (Lm7210 and Lm9252) and *L. turionifera* (Lt9434) had HCNTDs with monomers of 154/174/187 bp and 60/105 bp respectively and consistent HORs (Figure 3A). We aligned the three HCNTD repeat arrays in Lt9434 to see if they were related, which revealed that they were the same HCNTD with different levels of nucleotide identity and no obvious chromosome association (Figure S4). In all three of the *L. japonica* genomes (Lj7182, Lj8627 and Lj9421), both the *L. minor* and *L. turionifera* HCNTDs were found on their respective sub-genomes, consistent with these repeat arrays being specific to the parental lines (Figure 3A). In *L. minor* HCNTD monomers were within the typical size range (147-200) for centromeric satellite arrays (Figure 3A). In *L. turionifera* however, the small size of the monomeric repeat (60/105bp) leaves open the possibility that this *Lemna* species at least has holocentric chromosomes (Figures 3A, S4), while in *L. minor* repeat distribution was equally consistent with polycentric chromosomes ^27^. Chromatin contact mapping can reveal distinct interchromosomal associations typical of monocentric, polycentric and holocentric chromosomes, respectively ^27^. None of the species in this study had the stark, transverse centromeric clustering of Hi-C contacts typical of monocentric chromosomes (Figure S1C). It is therefore possible that at least some Lemnaceae spp. Are holocentric, though further experimentation is required. Exceptionally, the hybrid *L. japonica* contact map had a consistent pattern of interhomeolog associations extending along the length of each chromosome (Figures 3B, S4). Similar contact patterns have been reported between phased holocentric homologs in sedges ^28^, but not between homeologs of monocentric allopolyploids ^29,30^.

**Figure 3.**
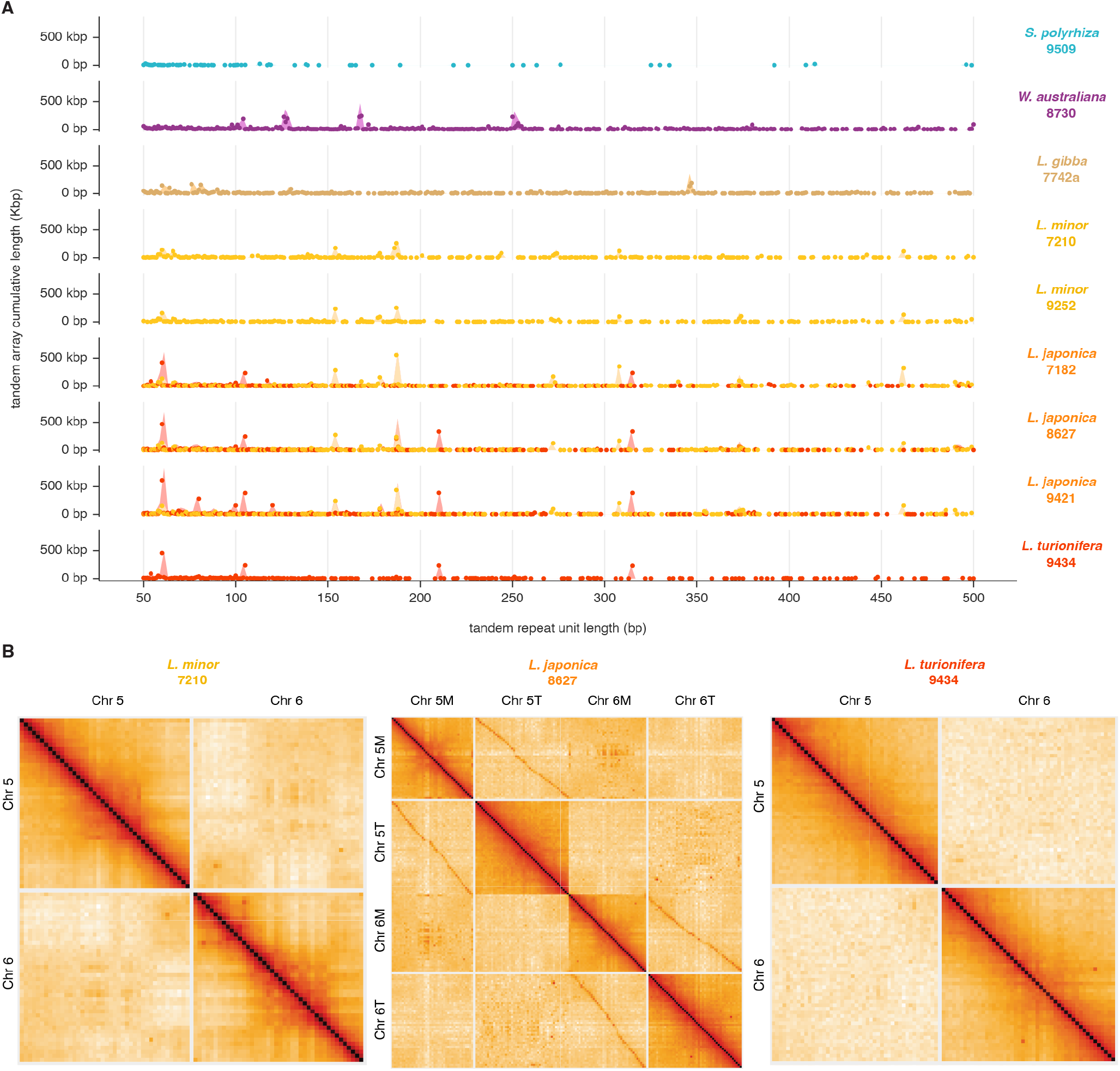
Satellite repeats are organized in limited tandem arrays resembling centromeres. **(A)** Tandem repeat arrays are plotted in each genome by repeat length (x axis) and cumulative array length (y axis). In *L. japonica* 8627 arrays have two distinct sets of base repeat units with higher order repeats (HOR) typical of centromeric satellites. The shorter 105 bp repeats are also found in *L. turionifera*, while the longer 154, 177 and 187 bp repeats are found in *L. minor* 7210. *Wolffia* has a distinct set of base repeats and HOR, while *Spirodela* lacks HOR. **(B)** Hi-C chromatin contact maps of chromosomes 5 and 6. Contacts are displayed at 500 Kbp resolution after balancing with the ICE method. None of the Lemnaceae chromosomes show clear centromeric contact clustering, and inter-homeolog contacts are evident in the *L. japonica* hybrid.

### Gene family gain and loss in the Lemnaceae

The genomes of the freshwater and marine aquatic plants *Spirodela polyrhiza*, *Wolffia australiana* and *Zostera marina* have a dramatically reduced gene set. For example, many genes for stomatal development are absent from *Zostera* ^31^ while genes for root development and disease resistance were lost in *Wolffia*. Consistent with a reduced morphology, single nucleus RNA-seq of the invasive *Lemna minuta* yielded a reduced molecular cell-type atlas ^19,26,32,33^. We undertook the first comprehensive multi-genera phylogenomic analysis of Lemnaceae together with other aquatic plants based on the proteomes of 11 Lemnaceae accessions (9 of which were annotated in this study), 15 additional angiosperms, and one gymnosperm outgroup. We determined that the divergence of the Lemnaceae occurred at the beginning of the Eocene, approximately 58 MYA (Figure 4A). We used OrthoFinder2 (Supplementary Methods) to infer phylogenetic relationships among the genes of these species (hierarchical orthogroups or “HOGs”), to discover common gene family losses and phylogenetically distinct paralogs across groupings of accessions (Figure 4B). To specifically examine adaptations to clonal reproduction and aquatic habits, we included *Ceratophyllum demersum*, a submerged, rootless freshwater coontail species considered to be sister to all eudicots, and *Zostera marina*, a monocot seagrass phylogenetically close to the Lemnaceae. Both species, like those in Lemnaceae, exhibit facultative asexual reproduction.

**Figure 4.**
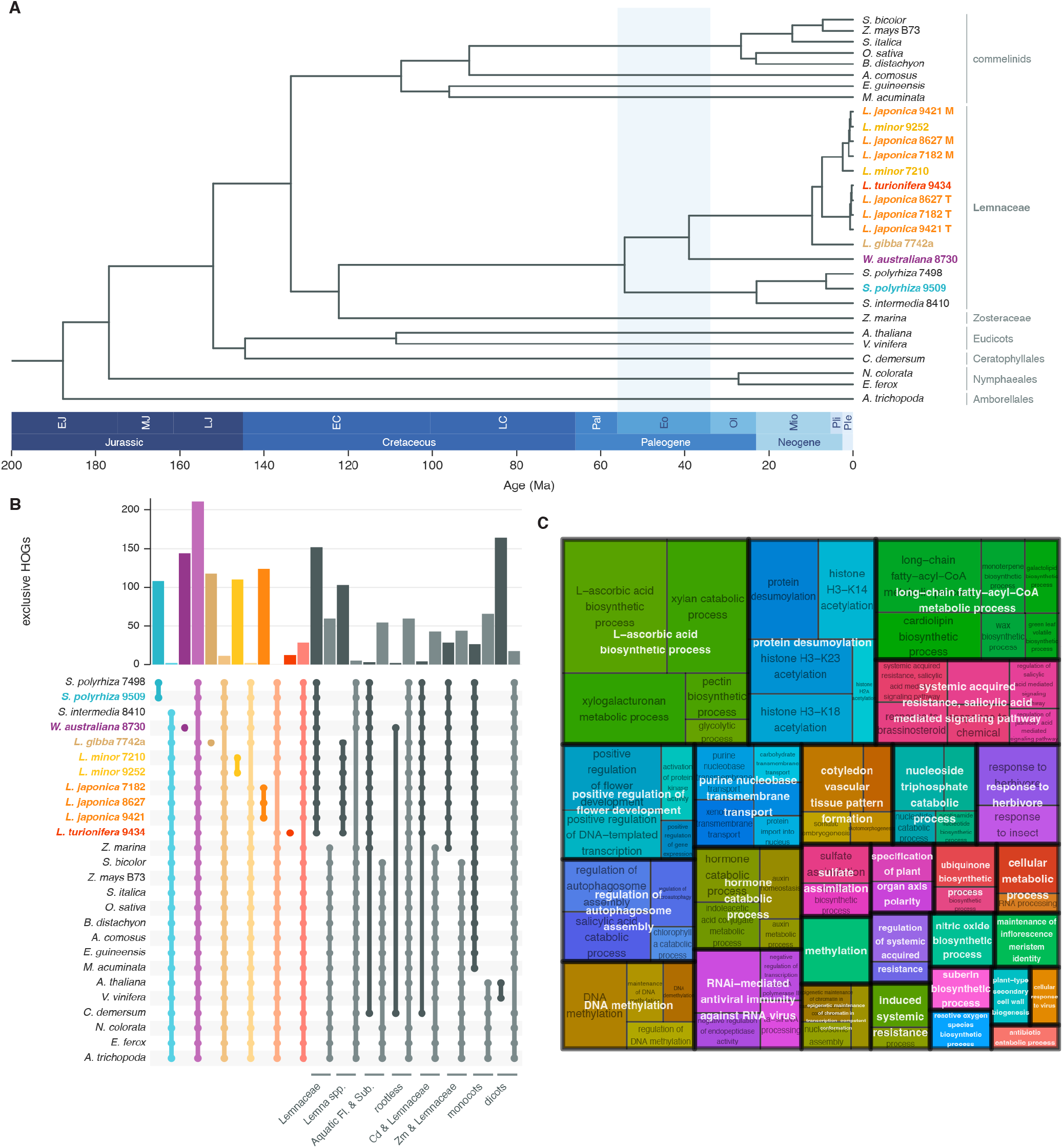
Phylogenomic analysis of the Lemnaceae. **(A)** Evolutionary relationship of chromosome-resolved Lemnaceae accessions to the angiosperms. The species tree topology was estimated from a concatenated supermatrix alignment (AA) of 854 genes identified as single-copy in 87% of the analyzed species, including 11 Lemnaceae clones, 16 other monocots and dicots, and *Gnetum montanum* (not shown) as the gymnosperm outgroup. Blue shading highlights the divergence of the Lemnaceae during the Eocene (Eo). **(B)** Unique paralogs and missing orthogroups. Groups of accessions from the full phylogenomic analysis are indicated by connected dots beneath counts of the common hierarchical orthogroups (HOGs) in each group. Groups are arranged in pairs, with the first showing unique paralogs, and the second showing missing HOGs present in all other accessions. Accessions with genomes annotated in this work are highlighted in color. Higher-order phylogenetic, phenotypic, and ecological groupings are represented by gray bars. Fl. & Sub. = floating and submerged. **c** Overrepresented GO terms in the set of HOGs missing from all Lemnaceae but present in both *A. thaliana* and *O. sativa*. GO terms were grouped by semantic similarity with the Relevance method and reduced with a cutoff of 0.7. Size of the rectangles is proportional to -log10(p-value) using Fisher’s exact test and a cutoff of 0.01.

This allows powerful predictions of genes that may be responsible for adaptations to clonal, aquatic and reproductive habits by grouping species that share these traits. Because of the unprecedented quality of single molecule genome assemblies, and consequently, highly accurate proteome prediction (Figure S5), we have been able to pinpoint gene and gene families including regulatory genes responsible for each adaptation. In total, we detected 60 missing HOGs in Lemnaceae, yet conserved in all other angiosperms, while 152 paralogous HOGs are found to be unique to this family (Figure 4B, Table S3). GO term analysis grouped predominant missing HOGs, and included genes required for flower and root development, organ polarity, stomatal closure and metabolic traits, and the striking loss of genes required for DNA methylation and RNA interference relative to the functionally annotated genomes of rice and Arabidopsis (Figure 4C).

#### Reduced morphology and growth habit

Lemnaceae lack root hairs and lateral roots, due to the absence of pericycle ^34^, yet we found that Lemnaceae do possess orthologs of key root development genes recently reported to be lost in *S. polyrhiza* ^35^. Namely, the Os*ZFP,* Os*NAL2/3* (*WOX3*), Os*ORC3*, Os*SLL1*, and Os*SNDP* families were present in all duckweeds. However, consistent with Wang et al.^32^, we found that all Lemnaceae have lost the root hair specific expansins At*EXPA7* and *18*, along with At*MYB93*, a very-long chain fatty acid responsive transcriptional regulator of lateral root development genes ^36^. *AtCMI1*, a Ca^2+^ sensor that regulates auxin response during primary root development ^37^, and *XAL2*, a transcription factor required for root stem cell and meristem patterning ^38^, were also absent in all duckweeds. *W. australiana* is rootless, and lacks *WOX5*, as previously reported ^26,39^, which encodes the homeobox transcription factor required for genesis of the meristem initials to start primary root development ^26^. This absence was shared exclusively with the other rootless plant in this study, *C. dememersum*, along with 59 other orthogroups (Figure 4B, Table S3). These include numerous root development genes, also missing from rootless carnivorous and parasitic plants ^40,41^ (*ARF5*, *RHD6*, *RGI1* and *2*, *DOT5* and *URP7*) (Figure 5).

**Figure 5.**
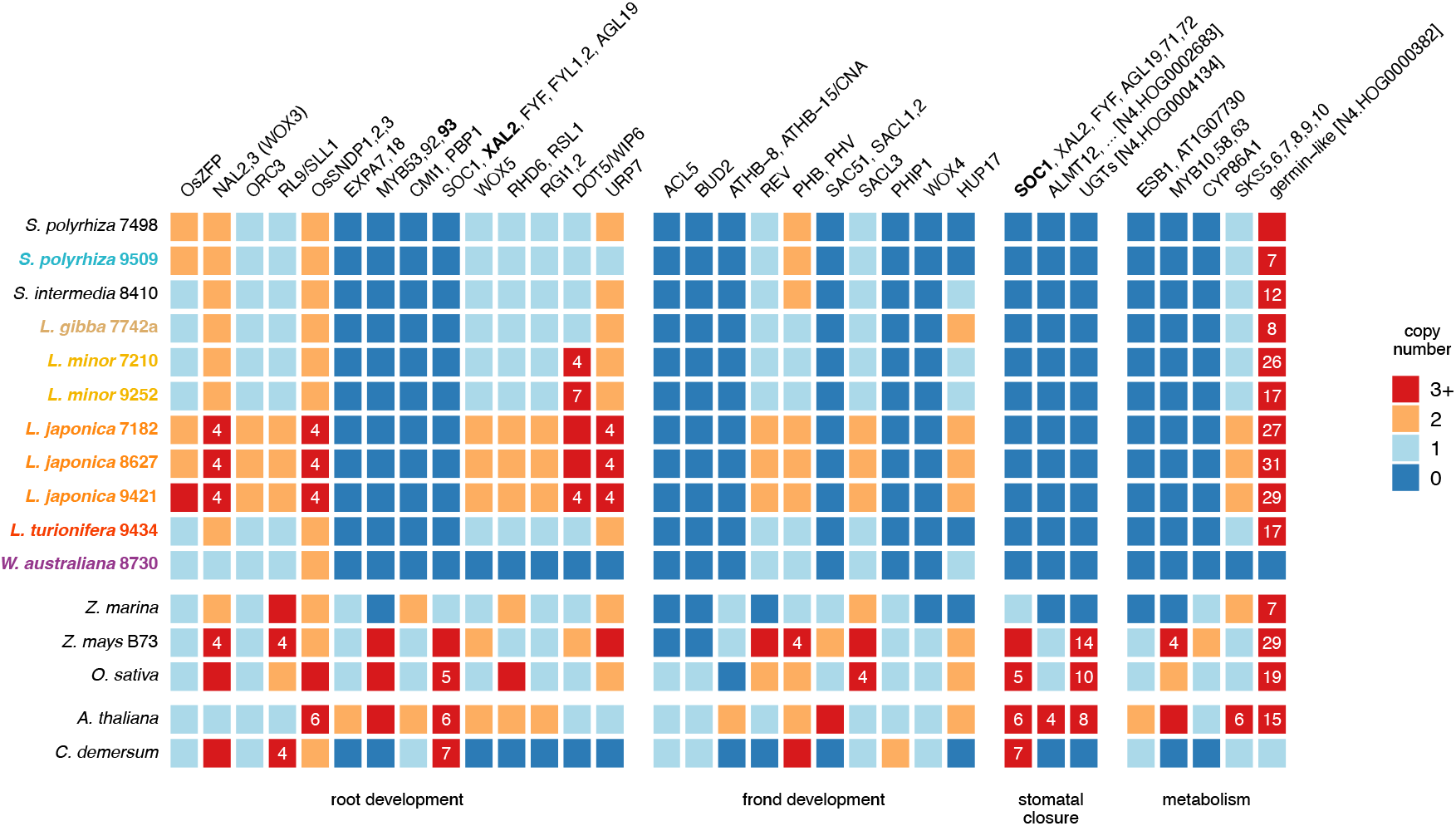
Gene losses shape Lemnaceae development, physiology, and metabolism. Selected gene families (hierarchical orthologous groups) were compared between the Lemnaceae, and submerged marine (*Z. marina*) and freshwater (*C. demersum*) aquatic plants, and compared to *Arabidopsis*, maize, and rice (Fig.4). Gene copy numbers are shown for selected genes involved in root development, frond development (stature, vascular patterning, polarity and turion formation), stomatal closure, and metabolism. Copy numbers from 0-3 are color coded as shown, higher copy numbers are indicated.

Turions are dormant buds induced by cold temperatures and low phosphate, and are found in many duckweed species including *S. polyrhiza* and *L. turionifera* (Figures 1A, 2A) ^42^ and *W. australiana*. In contrast, *L. minor* does not form turions, prompting us to examine whether this trait was retained in hybrids. We assayed turion induction in multiple accessions of *L. minor, L. turionifera* and their hybrid *L. japonica*, including Lm9252 which was phylogenetically closer to the M subgenomes of *L. japonica* hybrids than Lm7210 from South Africa (Figure 4A). Although growth rates were comparable, reaching a maximum of 24 mg per mg starting weight after only 7 days, we found that while *L.turionifera* readily formed turions under inductive conditions, 5 different accessions of *L. minor* and 6 of *L. japonica* did not form turions (Table S4). One interpretation is that *L. minor* has a dominant inhibitor of turion formation missing from *L. turionifera* but retained in *L. japonica*. One candidate is *HUP17,* a gene induced by hypoxia in *Arabidopsis* that promotes senescence after prolonged submergence ^43^. The ortholog in *L. minor* was missing from *L. turionifera* but present in both subgenomes of *L. japonica*. The loss of HUP17 might contribute to the endurance of turions, whose high starch content and contracted intercellular air spaces promote submergence in winter. HUP17 was also missing from *S. polyrhiza* and the other turion-producing plants in this study (*Z. marina*, *C. demersum*), with the exception of *W. australiana* 8730, an isolate from subtropical New South Wales which is unlikely to experience prolonged winters (Figure 5).

A broadly conserved, thermospermine-mediated tissue development regulatory module consisting of ACL5, BUD2, and the *HD–ZIP III* family transcription factor ATHB-8 was found to be absent from Lemnaceae. Loss-of-function mutants of the thermospermine synthase gene *ACL5* and the S-adenosyl-methionine decarboxylase gene *BUD2* exhibit severe dwarfism in *Arabidopsis*, along with xylem overproliferation and defects in auxin transport influencing vein development. Mutants of their upstream transcription factor *ATHB-8* disrupt the formation of the preprocambium and procambium, as well as xylem specification and differentiation ^44^. The absence of *ACL5*, which is variable in the monocots ^45^, has been observed previously in *S. polyrhiza* ^46^, and here we found that all Lemnaceae additionally lack *BUD2*, *ATHB-8*, and *CORONA*. These latter two genes antagonize the roles of the other *HD-ZIP III* members *REVOLUTA* (*REV*), *PHABULOSA* (*PHB*), and *PHAVULUTA* (*PHV*) in meristem formation, organ polarity, and vascular development ^47^. Duckweeds retain *REV*, but have just one homolog of *PHB* or *PHV*, while at least two paralogs were found in all but one other species. The bHLH transcription factors SAC51, SACL1, and SACL2 that participate in the ACL5-auxin feedback loop are also absent ^48^. While *SACL3* is retained, *PHIP1* ^49^ is lost, which could enhance *acl5* dwarfism, producing the “tiny-plant” phenotype found in *acl5 sacl3* mutants of Arabidopsis ^48^. *WOX4* is also absent, which regulates cell division in the procambium ^50^, and together with downstream factors such as *ATHB-8,* likely contributes to the dramatic simplification of the vascular bundle in Lemnaceae ^34^. Anatomical reduction, diminished vasculature, and altered leaf polarity (e.g. the presence of adaxial, rather than abaxial stomata supportive of gas exchange in a floating habitat) could be accounted for by these losses^47^ (Figure 5).

#### Stomatal response to elevated atmospheric carbon

We found that the key flowering regulator *SUPPRESSOR OF CONSTANS 1 (SOC1)* was missing from all analyzed Lemnaceae (N4.HOG0006359), contrary to a recent study in the short day duckweed *L. aequinoctialis* ^51^, but in accord with prior analysis of MADS-box genes in *S. polyrhiza* ^32^. In *Arabidopsis*, *SOC1* controls drought induced flowering ^52^ as well as light induced stomatal opening ^53^. Neither function would be required in duckweed fronds, which typically have open stomata and are not subject to drought. The *SOC1* paralogs *XAL2*, *FYF*, and *FYF1*,*2* are also missing, and impact various aspects of root and floral development ^38,54^. Lemnaceae, *Z. marina*, and *C. demersum* also lack orthologs of the guard-cell expressed aluminum-activated malate transporter *ALMT12*, which is largely responsible for the stomatal closure response during drought stress, and also involved in the closure response to CO_2 55,56_. A high copy family of UDP-glycosyltransferases (UGTs, N4.HOG0004134) involved in defense response accounted for the significantly enriched GO term “abscisic acid-activated signaling pathway involved in stomatal movement” was also absent from all duckweeds, *C. demersum*, and *Z. marina* (Figure 5).

#### Regulation of metabolic pathways

Duckweeds have simplified metabolic pathways that are reflected in both missing and uniquely paralogous orthogroups in each species. This is particularly true of polyphenolic metabolism responsible for structural rigidity of cell walls in terrestrial plants. One interesting example is the architecture of the Casparian strip, a lignified cell wall important for water transport. The Casparian strip (CS) has been observed in duckweeds, but has substantially reduced lignin content ^34^. The complete absence of Dirigent protein ESB1 responsible for lignin and alkaloid biosynthesis, and for organization of the CS, is consistent with this observation. The transcription factors *MYB58* and *MYB63*, which activate lignin biosynthesis during secondary cell wall formation ^57^ are also missing. *Lemna* and *Wolffia* have drastically reduced xylem and lack a defined shoot endodermis, likely reflecting this loss ^58^. The lipid biopolymer suberin is thought to form a diffusion barrier for water, gasses and solutes in lamellae that surround the CS, as well as in roots, where its engineered overproduction has been proposed as an inert polymer carbon sink for carbon sequestration applications ^59^. The cytochrome P450 monooxygenase *CYP86A1* is important for cutin biosynthesis in roots and seeds ^60^, and was found to be missing only in Lemnaceae and the rootless *C. demersum*. In another example, most duckweeds accumulate calcium oxalate crystals in calcium rich media, which sequester CO_2 61_ but could be problematic for mammalian consumption. Radiolabeling studies in *L. minor* and other plant species have demonstrated that ascorbic acid is likely to be the predominant source of oxalic acid that gives rise to crystals sequestered in idioblast cells ^62–64^. *Wolffia* is an exception, making neither druses nor raphides, and we found that *Wolffia* specifically lacks the *SKS5*-*8* ʟ- ascorbate oxidase orthogroup (N4.HOG0002231), providing a possible explanation as well as a target for genetic modification. A large family (N4.HOG0000382) of germin-like proteins possessing an oxalate oxidase enzymatic domain is also missing only in *Wolffia*, consistent with the loss of this substrate (Figure 5).

#### RNA interference, DNA methylation and gene silencing

The *Spirodela polyrhiza* genome has one of the lowest levels of DNA methylation found in any angiosperm ^19^. Low methylation levels could be a feature of clonal reproduction as DNA methylation levels in flowering plants are typically reset in the embryo ^65^ and are lost during somaclonal propagation ^66^. We therefore performed whole genome bisulfite sequencing of *L. gibba*, *L. japonica*, and *W. australiana* to determine if methylation loss was shared with *S. polyrhiza* (Figure 6A). We also sequenced small RNA from vegetative fronds from each of the four species (Figure 6B). We profiled coverage in both datasets over protein coding regions and interspersed repeats including LTR retrotransposons (LTR RTs) and DNA transposable elements (TEs) (Figure 6C).

**Figure 6.**
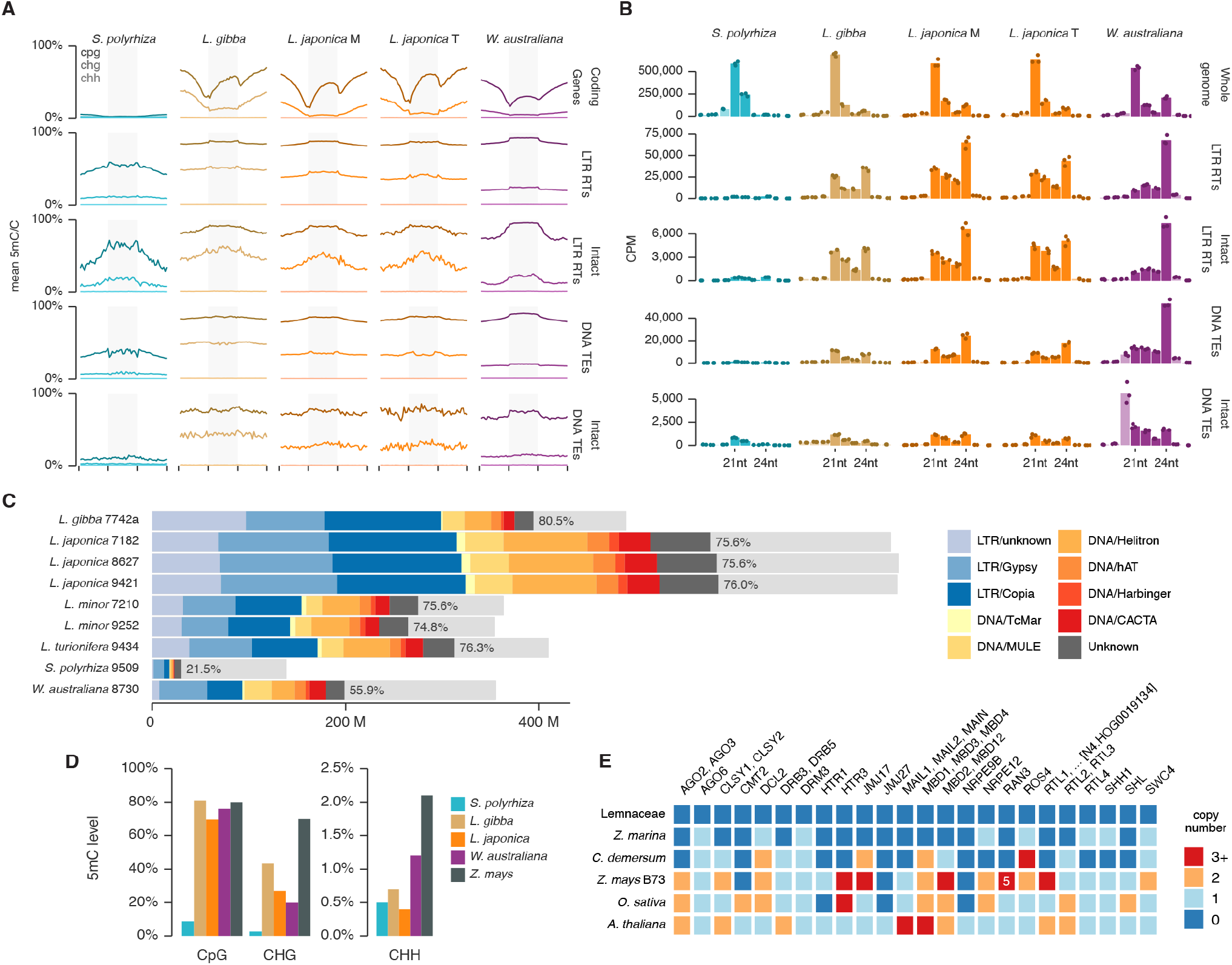
DNA methylation and small RNA regulation in the Lemnaceae. **(A)** Whole genome bisulfite sequencing (WGBS) of Sp9509, Lg7742a, Lj8627 and Wa8730 was used to generate combined metaplots of cytosine methylation levels in the CpG, CHG and CHH contexts over annotated regions (light grey bars) and 2 Kbp upstream and downstream windows. Regions shown: coding genes, LTR retrotransposons (LTR RTs), intact LTR retrotransposons (recently transposed), DNA transposable elements (TEs) and intact TEs, defined by intact ORFs and terminal inverted repeats (TIR). **(B)** Small RNA sequencing from fronds of each species was used to generate size distribution plots of 18-26nt small RNA, that mapped to the whole genome, or to the transposon classes shown in part **a**. Measurements from biological replicates (n=3) are plotted as points, and their means, as bars. 21-24nt small RNA are plotted in a darker shade. **(C)** Transposable element repeat family content of each genome, color coded as shown, and expressed as total length repeat-masked for each family in Mbp. Light gray bars show the size of each genome. **(D)** Global cytosine methylation levels in each sequence context as determined by WGBS. **e** Hierarchical ortholog groups (HOGs) for small RNA and DNA methylation gene families missing in the Lemnaceae, and compared with aquatic plants, maize (*Z. mays* B73), rice and *Arabidopsis*. Color coding as in Figure 5.

We found that all the duckweed genomes had low genome-wide levels of CHH methylation (0.4-1.2%), even less than in maize (2%) ^67^ and close to the limit of detection. However, levels of methylation were much higher in the other Lemnaceae at CG (69-81%) and CHG (27-43%) sites, compared to *S. polyrhiza* (8.9% CG and 2.7% CHG) (Figure 6D). Significant levels of CG methylation were detected in all four duckweed genomes, including *Spirodela* (Figure 6A). This indicates that the low level of CG methylation found in *S. polyrhiza* reflects the lack of intact transposons in this species. Strikingly, CG methylation in gene bodies was absent from *S. polyrhiza*, when compared with the other *Lemnaceae* (Figure 6A). Absence of gene body methylation in angiosperms is thought to be an indirect consequence of the loss of CHG methylation, even though CHG methylation is restricted to transposons ^68^. Consistently, CHG methylation is absent from *S. polyrhiza* transposons and reduced in *Wolffia*, which also has reduced gene body methylation (Figure 6A). Small RNA sequencing revealed a predominance of 21nt miRNA over 24nt siRNA compared to other angiosperms when mapped to the whole genome (Figure 6B). When mapped to transposons, however, we found that *S. polyrhiza* had very low levels of 24nt siRNA as previously reported ^69^, but the other species had much higher levels, corresponding more or less to the number of TEs in each genome (Figure 6C).

Next, we examined gene losses in the Lemnaceae that might account for these patterns of small RNA accumulation and DNA methylation. 24nt small RNA precursors depend on the RNA polymerase Pol IV, and the SWI2/SNF2 chromatin remodeler genes *CLSY1-4* along with the H3K9me2 reader *SHH1* are required for PolIV activity ^70^. HOG analysis revealed that all duckweeds have lost *CLSY1*, *CLSY2*, and *SHH1*, consistent with relatively low levels of 24nt sRNAs in vegetative fronds compared with other angiosperms (Figure 6E). However, retention of *CLSY3* in duckweeds suggests that 24nt siRNAs may be prevalent in the germline, where *CLSY3* regulates 24nt siRNA production in *Arabidopsis* ^70,71^. Small RNA are generated from precursors by Dicer-like RNAse III enzymes and loaded onto Argonaute RNAseH proteins that are required for stability and function of small RNA in silencing genes, transposons and viruses. We found that the Lemnaceae encode Dicer-like genes from only 3 of the 5 angiosperm clades, as in *Spirodela* ^19^. These Dicer-like genes are responsible for 21/22nt miRNA (*DCL1*), 21nt secondary small RNA (*DCL4*) and 24nt siRNA (*DCL3*), but all duckweeds lack *DCL2* and *DCL5*, which are responsible for 22nt and 24nt secondary small RNA, respectively. Duckweeds also have a reduced set of only 5 Argonaute genes, compared to 10 in *Arabidopsis*, 19 in rice and 22 in maize. They include Argonautes from each of the three major clades, which are predominantly associated with 21/22nt, and 24nt small RNA, respectively. This is consistent with small RNA sequencing from fronds that revealed all size classes of small RNA in each species, although the relative abundance varies with the abundance of different classes of transposons (Figure 6B). Intact DNA transposons in *Wolffia* have high levels of 20-21nt siRNA, consistent with transcriptional activity and post-transcriptional silencing ^72^, as do the very few DNA TE copies present in *Spirodela* (Figure 6B).

The Lemnaceae are specifically missing *AGO2*, *AGO3*, *AGO6* and *AGO9* compared to maize and *Arabidopsis*. These argonautes are highly expressed in *Arabidopsis* pollen and seeds, and the absence from the Lemnaceae could reflect their clonal growth habit ^65,73^. Similarly, *DCL5* is normally expressed in the male germline of many monocots, where it is required for fertility ^74^. *DCL2* is thought to be responsible for viral resistance in many angiosperms, and it is possible that Lemnaceae have less need for this particular antiviral strategy, although other aquatic plants have retained it. The lack of viral defense RNAse III-like genes *RTL1,2,* and *3* supports this conclusion. Intriguingly, however, the only other angiosperm in this comparison to lack *DCL2* is the African oil palm *E. guineensis*. This may reflect a role for DCL2 in reproductive isolation, as oil palm, like duckweed, is interfertile with distantly related species despite many millions of years of divergence ^75^. *DCL2* has recently been found to be responsible for hybrid incompatibility and meiotic drive in maize and its relatives ^76^.

DNA demethylation by ROS1 requires the histone H3K18 and H3K23 acetyltransferase gene *IDM1/ROS4* which encodes a conserved protein in the IDM complex that acts upstream of H2A.Z deposition by SWR1. In *Arabidopsis*, this mechanism rescues some euchromatic regions from promiscuous RdDM targeting ^77^. Among all angiosperms analyzed here, loss of *IDM1* is only observed in duckweeds. This could reflect the very low levels of CHH methylation in vegetative fronds, making demethylation unnecessary. However, the presence of *DRM1* and *2* suggests CHH methylation is still possible, most likely in seeds and pollen grains, which have the highest levels in *Arabidopsis*, but which were not examined here. Instead, the DNA methyltransferase gene *CMT2*, conserved in most other sequenced plant genomes, is responsible for high levels of heterochromatic CHH methylation in rice and *Arabidopsis*, but is absent from duckweeds and maize accounting in large part for the low levels of CHH methylation (Figure 6E) ^78^.

#### Reproduction and clonal growth habit

Polyploidy, aneuploidy, and mixoploidy have all been observed among the Lemnaceae ^79^, and a recent meta-study of chromosome counts and genome size estimates indicate that triploidy occurs frequently in *Lemna* and *Wolffia*, with 2n ≈ 60 appearing in 9 of the 36 currently recognized Lemnaceae species, including *L. japonica* (*L. minor* x *L. turionifera*) ^80^. Plastome-based barcoding has demonstrated that *L. japonica* hybrids are always formed from an *L. minor* seed parent ^9^, assuming maternal inheritance of plastids. Sequence identity comparisons of the *de novo* assembled plastid and mitochondrial genomes in this study support this conclusion (Figure S6). Furthermore, the absence of RanGAP (N4.HOG0008339) genes in Lt9434 and the Lj9421 T subgenome suggests that at least some *L. turionifera* lineages might not produce viable female gametes ^81^. The two major paths to polyploidy in angiosperms are somatic doubling and gametic non-reduction, with the latter being a more frequent contributor ^82^. Non-reduced gamete formation resulting from abnormalities in both micro-and megasporogenesis is heritable, and much more frequent in hybrids ^83^. Since viable triploids with diploid contributions of either parental genome are possible (Lj7182 and Lj8627) and heterozygosity is evident in both cases (Figure 2C), unreduced gamete formation in both the *L. minor* maternal and *L. turionifera* paternal germlines is a likely explanation for the emergence of these interspecific hybrids.

Unreduced maternal gametes arise via diplospory in maize mutants of *ago104*, the ortholog of *AGO9* in *Arabidopsis*, and retain heterozygosity in unreduced gametes and their progeny ^84^. *ago9* mutants in *Arabidopsis* also have the potential to form diploid gametes via apospory, as they produce supernumerary megaspore mother cells that differentiate directly from diploid somatic cells ^85^. The Lemnaceae only have one paralog in the *AGO4, AGO6, AGO8, AGO9* clade, which appears to be related to *AGO4*. *AGO9* is highly conserved among angiosperms, and its loss from duckweeds is an unusual feature that could account for the origin of triploid hybrids such as *L. japonica* 8627, whose maternal *L. minor* parent appears to have had unreduced heterozygous gametes. In contrast, *L. japonica* 7182 has two copies of the paternal *L. turionifera* genome, indicating unreduced paternal gametes. So far, direct observations of pollen development in Lemnaceae have been limited to *L. aequinoctialis* (formerly *L. paucicostata* HEGELM.) ^86^, which was found to be tricellular. One candidate explanation for the production of 2n male gametes is disruption of *JASON* (*JAS*), a positive transcriptional regulator of *PARALLEL SPINDLES1 (AtPS1)* in meiotic cells which is required for pollen meiosis II spindle polarity, but not involved in female meiosis. In *Arabidopsis*, homozygous mutations in *JAS*, as in *AtPS1*, cause heterozygous 2n pollen formation at rates up to 60% ^87–89^ and result in fertile haploids ^90^. We found deletions and mutations in each of the two *JAS*-like loci in *Lemna* spp. in regions deeply conserved across other taxa (Figure S7A). None of the predicted *JAS* orthologs in aquatic plants possess the N-terminal Golgi localization peptide found in terrestrial plants under hypoxia ^91^.

Triploid hybrids are rare among angiosperms, due to the “triploid block”, a prevalent form of reproductive isolation in which seeds fertilized by unreduced diploid pollen abort. The triploid block depends on the level of small interfering RNA in pollen, and mutants in several genes in the RdDM pathway reduce or eliminate the triploid block in *Arabidopsis* ^10,92^. These mutants include *ago6*, which is absent from the Lemnaceae, but completely conserved in other taxa (Figures 4C, 6D), potentially accounting for the prevalence of triploids. Sexual reproduction strongly selects against triploids due to aneuploid swarms, whereby unequal segregation in triploid meiosis results in aneuploidy and severe fitness penalties ^93,94^. Clonal reproduction from germinating triploid seeds avoids meiosis and enables other advantages of increased heterozygosity and gene dosage.

*MSH4* and its heterodimer partner *MSH5* (MutSγ) are meiosis-specific mismatch repair proteins in the ZMM pathway required for the formation of Class I crossovers responsible for 80% to 90% of chiasmata via stabilization of double Holliday junctions ^95,96^. After polyploidization in plants, meiotic recombination genes are the most rapidly fractionated in the genome ^97^, and while supernumerary MutSγ copies do not increase total crossover number, reduction to a single copy per subgenome prevents inter-homeolog crossovers, benefitting chromosome segregation in hybrids ^98^. The *MSH4* orthologs in Lm9252 and the M subgenomes of Lj8627 and Lj9421 share a 163 residue N-terminal truncation, entirely eliminating the Holliday junction-binding MutSII domain. In the homozygous Lm7210, this extends to 234 residues, partially encroaching on the MutSIII domain (Figure S7B). Similar N-terminal truncations of *TaMSH4D* and *TaMSH5B* resulting in pseudogenization have been noted in the subgenomes of allohexaploid wheat and its ancestral tetraploids ^99^. In contrast, all other Lemnaceae orthologs, including that of Lj7182 (MTT) subgenome M, are full length.

## Discussion

The chromosome-level genome sequence assemblies reported here provide insight into the evolutionary history, reduced morphology, and reproductive growth habit of the Lemnaceae, the world’s smallest but fastest growing flowering plants. Our evolutionary analysis suggests that the Lemnaceae arose in the Cretaceous but diverged in the mid-Eocene, coincident with the “Azolla event”, when arctic core samples suggest that huge blooms of the freshwater aquatic fern Azolla grew in the inland palaearctic sea ^100^. These blooms are thought to be responsible for 90% reduction in atmospheric carbon, from 3600 to 300 ppm, in less than a million years. Although they are much harder to detect in core samples, Lemnaceae fossils have been found in shale deposits from this time and likely cohabited these warm freshwater environments ^101^. Aquatic plants are uniquely adapted to high CO_2_ environments, as stomatal closure in response to elevated CO_2_ has been lost in many species, and photosynthetic rate can thus increase with rising CO_2_. We found that, unlike the submerged seagrass family *Zostera*, Lemnaceae have retained key patterning genes required for guard cell formation, but they have lost at least 3 master regulators of stomatal closure in response to light, drought and CO_2._

Along with genes required for lateral roots and root hairs, we found that genes for acquired and systemic disease resistance are largely missing from *Lemna* as in *Wolffia* and *Spirodela* ^19,26^. This likely reflects adaptive advantage in their common floating freshwater habitat shared with waterfowl and other metazoans. In addition, a wealth of genes encoding and regulating metabolic enzymes are either missing, or have unique paralogs in *Lemna*. Examples include unique and missing paralogs in long chain fatty acid biosynthesis, and in the suberin biosynthetic pathway. We have recently engineered *L. japonica* to produce and accumulate 100 times more oil (triacylglycerol) than in wild type fronds, and long chain lengths in this context are consistent with our findings ^13^. Suberin accumulation in roots has been proposed as a strategy for carbon sequestration in terrestrial crop plants ^59^ but would need engineering (like oil) to be successful in Lemnaceae. An alternative carbon sink could be calcium oxalate, and we identify the biosynthetic pathway found in duckweeds. Finally, Lemnaceae are a promising high protein crop, in part due to reduced cell size relative to the number of plastids ^1^, which provide most of the protein in leaves. We identify the loss of a spermine-TF module that is likely responsible for the reduced stature and adaxialized polarity which underlie this key trait.

The clonal growth habit of Lemnaceae and other aquatic plants has resulted in dramatic changes in chromosome biology and epigenetic regulation, consistent with prolonged clonal expansion in the absence of meiosis. Loss of transposons is thought to be a consequence of clonal growth habit, as transposons require meiotic recombination to increase in copy number ^102–104^. The more drastic loss of transposons in *Spirodela* in comparison to *Lemna* and *Wolffia* may reflect decreased propensity for sexual reproduction in *Spirodela* ^17,105^. Consistently, *Lemna* and *Wolffia* have far greater numbers of recently active LTR retrotransposons as evidenced by high identity LTR sequences and high levels of CpG methylation. The Lemnaceae have lost several genes encoding key components of the RdDM pathway, notably *CLSY1* and *2*, as well as *AGO6* and *AGO9*. *AGO6* is responsible for de novo transgene silencing ^106^ making duckweed potentially more permissive for transgenic applications. But why would it be advantageous for a clonally propagated plant to lose this aspect of gene silencing? Ectopic DNA methylation occurs spontaneously in seed plants, and depends on RdDM ^107^, but is reprogrammed during reproductive development, which removes epigenetic variation in pollen, and re-establishes parental patterns of methylation in the seed ^65,108^. Clonally propagated Lemnaceae do not undergo meiosis, and therefore do not undergo reprogramming, for thousands of asexual generations at a time, potentially leading to clonally inherited deleterious epigenetic variation ^109^. Therefore, losing at least some aspects of *de novo* methylation would mitigate these risks.

The downside of losing *AGO6* and other components of RdDM is that duckweeds appear to have lost the triploid block, allowing the formation of triploid hybrids when reproduction does occur. But triploids are only problematic for sexual, not clonal, reproduction, and clonally dividing cell cultures, at least in *Arabidopsis*, also dispense with RdDM ^66^. It is quite possible that some level of RdDM might be restored in seed and pollen, when cell division ceases, as observed in *Arabidopsis* ^65,66^. One explanation for the prevalence of polyploidy in the Lemnaceae is the presence of apparently defective homologs of *JASON*, that could result in high frequencies of unreduced paternal gametes, and the loss of *ago9* which could result in unreduced maternal gametes. Finally, defective orthologs of *MSH4* in the Lemnaceae would reduce homeologous recombination in balanced polyploids, promoting fertility as in other polyploid species ^110^. The presence of inter-homeolog contacts in *L. japonica* Hi-C maps suggests that recombination could be potentiated if these associations also exist in meiotic cells.

Mapping of putative centromeric satellite repeats in the Lemnaceae indicates that while some species appear to have well defined higher order tandem repeats typical of centromeric satellites, *L. turionifera* has much shorter satellite repeats that are not obviously associated with each chromosome. Our Lemnaceae Hi-C maps all lack the transverse contact pattern found in monocentric species at a single locus per chromosome, and instead more closely resemble the recently characterized holocentric chromosomes in sedges ^27^. This raises the possibility that some, if not all, species may be polycentric or holocentric. Hybrids between monocentric, polycentric and holocentric species have not previously been reported ^111^, and could potentially lead to genome instability. Orthogroup analysis revealed that all maternal subgenomes of *L. japonica* hybrids have lost the centromere cohesion factor shugoshin (*SGO1*)^112^. *SGO1* may play a role in the promotion of inverted meiosis by which holocentricity can overcome reproductive barriers that arise through chromosomal rearrangements ^111^. Intriguingly the only such rearrangement we detected in *Lemna* was between chromosome 17 in *L. minor* and chromosome 20 in *L. turionifera*, accompanied by a new rDNA locus. In this case, holocentricity is predicted to retain fertility in both the *L. turionifera* parent and the hybrid *L. japonica*.

In summary the complete genomes of clonal aquatic macrophytes, including both submerged seagrass and the floating Lemnaceae, pave the way for understanding and exploiting their regular division as novel crops, robust platforms for biomass and biotechnology applications, as well as their ancient and enormous potential for climate amelioration.

Draft versions of the genome assemblies and annotations presented here were released ahead of publication at www.lemna.org ^113^ and have already been utilized in several studies ^4,5,15,114–117^.

## Methods

### Sample preparation

Sterile cultures of the 9 accessions in Figure 1 were obtained from the Rutgers Duckweed Stock Cooperative and cultivated in 50 mL Schenk and Hildebrandt (SH) medium with 1% sucrose at pH 5.6 ^118^ at 23°C under a 16 hour photoperiod of approximately 30 μmol/m2/s per second of white fluorescent light. For HMW DNA extraction, culture flasks were covered in foil and grown in the dark for up to 2 weeks prior to harvest to deplete excess carbohydrates. Cultures were then flash frozen in liquid N_2_ and stored at -80°C.

### HMW DNA extraction

High molecular weight (HMW) DNA extractions were performed for Lg7742a, Lj8627, and Wa8730 using a modified CTAB prep followed by a high-salt low-ethanol starch cleanup (Pacific Biosciences) as described previously ^76^, except that 4 g of frozen duckweed tissue was used and the sorbitol wash was omitted. HMW DNA was isolated for Lm7210 using a modified Bomb protocol as described (https://bomb.bio/protocols/). HMW DNA was isolated for Lm9252 and Lj7182 using a modified version CTAB/PVP protocol as previously described ^119^. For Lj9421 and Lt9434, a modified nuclear extraction was used as previously described ^120^.

### Oxford Nanopore library preparation and sequencing

Long-read data for Lg7742a, Lj8627, and Wa8730 were collected over several years as the MinION sequencing platform matured. Both the SQK-LSK108 and SQK-LSK109 kits were used to prepare libraries. The method used to produce the most recent runs in this study is described in^76^. Completed libraries were loaded onto R9 or R9.4.1 flow cells and sequenced on the MinION instrument. Libraries for Lj7182, Lj9421, Lm7210, Lm9252, and Lt9434 were prepared as previously described ^33^ and sequenced on R9.4.1 flow cells on the PromethION platform using the rapid sequencing barcoding kit (SQK-RBK004). Offline base calling of all ONT reads was performed with Guppy 5.0.7 and the R9.4.1 450bps SUP model on NVIDIA Tesla V100 GPUs.

### Short read whole genome sequencing

Short read WGS libraries for Lg7742a and Lj8627 were prepared from 2µg of HMW gDNA using the Illumina TruSeq DNA PCR-Free kit (Illumina, cat#20015962) and sequenced on an Illumina MiSeq (PE 250bp, PE 300bp) or HiSeq 2500 instrument (PE 150bp). Libraries for other *Lemna* accessions were prepared as described for the NCBI SRA experiment SRX7624904, and previously published libraries were used for Wa8730 (SRX8008794, SRX8008795) ^26^. Illumina gDNA reads were aligned to the reference assemblies with bwa-mem2 v2.2.1 and coverage was calculated over 1bp bins with deeptools v3.5.2 ^121^ “bamCoverage --samFlagExclude 2304 -- ignoreDuplicates --binSize 1 --normalizeUsing CPM”.

### HiC library construction and sequencing

For Lg7742a, Lj8627 and Wa8730, approximately 10g of tissue per sample was sent to Dovetail Genomics and Hi-C libraries were prepared using their in-house protocol with the DpnII enzyme. For Lm7210 and Lt9434, the Phase Genomics Proximo HiC Kit (Plant) was used to prepare libraries^33^. The resulting libraries were sequenced on an Illumina NextSeq 500 (PE 150bp). The Juicer pipeline v2.20 ^122^ with options “-s DpnII” was used to align reads back to the final assemblies and construct initial contact maps (MAPQ ≥ 30) before multi-resolution cool file conversion with hic-straw v0.0.8 ^122^, ICE balancing with Cooler v0.9.2 ^123^, and visualization with HiGlass v1.11 ^124^.

### Heterozygous variant calling

ONT reads were aligned to their respective reference assemblies with minimap2 v2.24-r1122 ^125^ “-x map-ont -- MD”. Single nucleotide variants and indels shorter than 50 bp (SNVs) were called using Clair3 v0.1-r12 ^126^ with options “--min_contig_size=1000000 --platform ont --model_path /opt/models/r941_prom_sup_g5014”. Structural variants (SVs) were called with Sniffles2 v2.0.7 with default options. SNVs and SVs were filtered using RTG Tools v3.12.1 (https://github.com/RealTimeGenomics/rtg-tools) with options “vcffilter --min-read-depth=10 --min-genotype-quality=20”, and homozygous calls were removed with “vcffilter --remove-hom”.

### Whole genome bisulfite sequencing and analysis

WGBS libraries were prepared for Sp9509 and Wa8730 as previously described ^19^. Lg7742a and Lj8627 were prepared as previously described ^65^ for *Arabidopsis* embryos, except that DNA from the HMW extraction methods described above was used. Libraries were sequenced on a NextSeq 500 (PE 151) or a HiSeq 2500 (PE 108). Adapter sequences were removed and reads were hard-trimmed with Trimmomatic v0.35 ^127^ with options “HEADCROP:5 TRAILING:3 MINLEN:25”. Technical and biological replicates were merged, and reads were aligned to the genomes using Bismark v0.23.1 ^128^ with options “-N 1 -L 20 --maxins 1200”. Reads were deduplicated and methylated cytosines were called using “bismark_methylation_extractor --CX --bedGraph -- ignore_r2 2 --comprehensive” and a genome-wide cytosine report was generated with coverage2cytosine, and separate bigWig coverage tracks were derived for CpG, CHG, and CHH contexts. Profiles over genomic regions were calculated with deepTools 3.5.1 ^121^ computeMatrix with options “scale-regions --skipZeros -bs 100 -m 2000 -b 2000 -a 2000” and plotted in R. Genome-wide methylation levels were determined by calculating the weighted methylation level (#C/(#C+#T)) ^129^ for nuclear genome cytosine positions with a coverage of at least 5 reads. For reference, approximate genome-wide methylation levels for *Z. mays* B73 were derived from ^67^.

### Nanopore direct methylation analysis

Direct 5-methylcytosine modification calling in all contexts from the ONT WGS reads was performed with Megalodon v2.5.0 (https://github.com/nanoporetech/megalodon) with the dna_r9.4.1_450bps_modbases_5mc_hac model and Guppy v5.0.7 on NVIDIA Tesla V100 GPUs. Calls for Cs with fewer than 5 supporting reads were discarded, and the resulting bedMethyl files were split by cytosine context (CpG, CHG, CHH) using bedtools v2.30.0 ^130^ intersect. Fractional methylation calls at each cytosine were adjusted to a [0..1] scale, and the bedMethyl files were converted to bigWig format. Profiles over genomic regions were calculated as for the WGBS libraries.

### Transcriptome sequencing

To provide broad transcriptional evidence for annotation, RNA samples were collected from fronds of Lg7742a and Lj8627 grown under a diverse set of conditions: variable daylength, nutrient stress, temperature stress, high NaCl, high pH, UV damage, and exogenous hormone exposure. Samples from all conditions were pooled, polyA enriched or rRNA depleted, and strand-specific cDNA libraries were prepared and sequenced on Illumina and Oxford Nanopore Technologies instruments.

### Small RNA sequencing and analysis

RNA was extracted from 3 biological replicates of 100mg of frozen tissue of each accession using the Quick-DNA/RNA Miniprep kit (Zymo) following manufacturer’s instructions with the following modifications: frozen tissue was ground under LN_2_ with a mortar and pestle and resuspended in the Shield solution. It was refrozen in LN_2_, thawed, and treated with Proteinase K, and frozen again after the addition of lysis buffer. After thawing, samples were centrifuged at full speed, RT for 3 min. to remove debris, and the manufacturer’s protocol was followed as described. Afterwards, samples were DNase treated, enriched for RNAs 17-200 nt in length, and concentrated using the RNA Clean & Concentrator kit (Zymo). Libraries were prepared with the NEXTflex Small RNA-Seq Kit v3 (Perkin Elmer) following manufacturer instructions and sequenced on a NextSeq 500 (SE 75). Sequencing adapters were trimmed using skewer v0.2.2 ^131^ and 4nt randomer sequences were removed from both ends of each read with fastp v0.20.1 ^132^ with options “--trim_front1 4 --trim_tail1 4 --length_required 10 -- length_limit 35 --disable_adapter_trimming --trim_poly_g -q 20 --unqualified_percent_limit 10”. Reads mapping perfectly to annotated structural RNAs and organelles were removed, and those remaining were aligned to the genomes with ShortStack v4.0.0 ^133^ using options “--mincov 0.5 --mmap f --dn_mirna --knownRNAs <*miRBase 22.1 mature plant sequences*>”. Separate ShortStack runs with merged biological replicates were performed for microRNA discovery, and microRNAs identified by ShortStack were annotated by top BLAST hit of the mature sequence to miRBase 22.1 ^134^. Alignments were split by size class and bigWig coverage tracks computed with deepTools “bamCoverage --binSize 1 --normalizeUsing CPM”. Genomic region profiles were computed as described for the bisulfite libraries and plotted in R.

### Genome assembly

Reads longer than 1 Kbp were assembled into contigs with Flye 2.8.3-b1722 ^135^ with options “--extra-params max_bubble_length=2000000 -m 20000 --plasmids -t 48 --nano-raw”. The same reads were then aligned to the assembly using minimap2 2.20-r1061 ^125^, and these alignments were passed to the PEPPER-Margin-DeepVariant 0.4 pipeline ^136^ to polish the initial consensus with default options. To correct remaining SNVs and small indels, Illumina gDNA libraries were mapped to the long read polished consensus with bwa-mem2 2.2.1 ^137^ for further polishing with NextPolish 1.3.1 ^138^. To reduce occurrences of uncollapsed haplotigs and heterozygous overlaps in the assemblies, purge_dups 1.2.5 ^139^ was run with options “-a 80 -2”. For the hybrids, contigs were first assigned to either the *L. minor* or *L. turionifera* subgenomes by performing an initial pseudomolecule scaffolding with Hi-C reads as described below, followed by sequence similarity ranking of pseudomolecules using MegaBLAST 2.11.0+ ^140^ alignment of *L. minor* 5500 contigs ^141^ against the target scaffolds. Target contigs comprising each parental pseudomolecule set were then independently treated with purge_dups. Next, Hi-C reads (PE150) were mapped to the polished, heterozygosity-purged contigs with the Juicer pipeline v1.6 ^122^ UGER scripts with options “-s DpnII”. The resulting “merged_nodups.txt” alignments were passed to the 3D-DNA pipeline to iteratively order and orient the input contigs and correct misjoins ^142^. The initial automatic scaffolding was followed by manual review with JBAT ^143^. No Hi-C data were available for accessions Lj7182, Lj9421, and Lj9252, and instead pseudomolecules were constructed using RagTag 2.0.1 ^144^ correct and scaffold steps with default options and the final Lj8627 assembly as a reference. For all accessions, A final haplotype-aware short read polishing step was performed with Hapo-G 1.2 ^145^ using default options. Assembled pseudomolecules for Lm7210 and Wa8730 were named chr{1..N} according to length. All other *Lemna* pseudomolecules were oriented and named according to homology with Lm7210 chromosomes. Unintegrated contigs were screened for viral and bacterial contaminants by MegaBLAST search against assembled target accession pseudomolecules, organelles, and the NCBI nt ^146^ databases simultaneously.

### Organelle genome assembly and annotation

Reference plastid (CP) and mitochondrial (MT) genomes were downloaded from NCBI RefSeq (*Spirodela* MT: NC_017840.1; *Lemna* CP: NC_010109.1, *Wolffia* CP: NC_015899.1) ^32,147,148^ and 180° rotations were generated using SeqKit 2.2.0 ^149^ sliding “-C <*reference*> -s <*ref length / 2*> -W <*ref length*>”. ONT reads at least 40 Kbp long were aligned with minimap2 2.22 ^125^ to the original and rotated versions of CP and MT references simultaneously. Reads aligning to each reference were extracted for separate *de novo* assembly with Flye 2.8.3- b1722 ^135^ with options “-m 20000 --asm-coverage 100 --nano-raw” (CP genomes) or “-m 20000 --meta --nano-raw” (MT genomes). A single CP contig of length ±10 Kbp relative to the reference was assembled for all accessions, with ∼700-800x downsampled coverage. In the MT case where numerous contigs > 150 Kbp were assembled for each accession, only the contig with the highest coverage (∼13x-104x) was retained. PEPPER-Margin-DeepVariant and then NextPolish with downsampled short reads were used to polish the assemblies, which were then manually oriented and rotated for consistency with psbA(-), ndhF(-), ccsA(+) ^150^. Assemblies were compared by calculating average nucleotide identities (ANI) ^151^ and constructing dot-plots from pairwise genome alignments using nucmer v3.1 ^152^ and an R script (https://github.com/tpoorten/dotPlotly/blob/master/mummerCoordsDotPlotly.R).

Organelle genomes were annotated using the web application GeSeq v2.03 ^153^. The following non-default settings were used to annotate plastomes: circular; Sequence source = Plastid (land plants)/Mitochondrial; Annotation revision = Keep all annotations; HMMER profile search: [enabled]; tRNAscan-SE v2.0.7: [enabled]; Chloe v0.1.0: Annotate = CDS+tRNA+rRNA. Inverted repeats were annotated using self-pairwise genome alignment with nucmer. Gene annotations were manually reviewed. blatN annotations were used for rRNA genes and Chloe annotations were selected for tRNA genes. For protein-coding genes, Chloe annotations were used for most protein-coding genes but in cases where Chloe annotations did not result in correct open reading frames (proper start and end codon), then blatX or HMMER annotations with the correct open reading frames were selected. To ensure the correct annotation of the Wa8730 plastome, the Wa7733 plastome assembly (GenBank accession JN160605-1) was downloaded, annotated as above, and compared to the Wa8730 reference.

### Repetitive and non-coding sequence annotation

De novo repeat libraries were constructed for each accession using EDTA v1.9.6 ^154^ with options “--anno 1 –cds <SP7498 CDS sequences> --sensitive 1”. A softmasked version of each assembly was generated with the EDTA make_masked.pl script with options “-minlength 80”. Tandem repeats were identified in the genome assemblies using Tandem Repeats Finder v4.09 ^155^ with options “1 1 2 80 5 200 2000 -d -h”. The resulting *dat files were reformed and each repeat length was summarized to identify putative centromere and telomere arrays as described previously ^26^. Repeats of a specific length were summed and plotted as a function of repeat length revealing potential centromere arrays (Figure 3a). Ribosomal DNA loci were identified (along with other conserved non-coding genes) using Infernal v1.1.4 ^156^ cmscan with Rfam 14.1 ^157^ with options “-Z <EFFECTIVE genome size> -- cut_ga --rfam --nohmmonly”. Lower-scoring overlapping hits were removed. Exact occurrences of telomere sequence were identified on both strands using SeqKit v2.2.0 ^149^ locate with options “--bed --ignore-case -p TTTAGGG”.

### Gene prediction and annotation

Protein coding gene annotation for each accession was performed with a combination of *ab initio* gene prediction, RNA transcript evidence, homologous protein evidence, and comparative gene prediction approaches. Softmasked versions of the assemblies were used for all steps.

First, all available short-read (SR) cDNA evidence was aligned with HISAT2 v2.2.1 ^158^ with options “--dta --max-intronlen 10000 --rna-strandness <*orientation*>”. An initial *ab initio* prediction set was built with BRAKER v2.1.6 ^159^ and TSEBRA v1.0.3 ^160^, incorporating the HISAT2 alignments and plant protein evidence from OrthoDB v10 ^161^ (*ab_initio_preds*).

Alternative SR alignments were made with STAR v2.7.9a ^162^ with no reference annotation with options “-- twopassMode Basic --alignIntronMin 20 --alignIntronMax 10000 --alignMatesGapMax 10000 -- outFilterMismatchNmax 4 --outFilterIntronMotifs RemoveNoncanonical --outSAMstrandField intronMotif -- outFilterMultimapNmax 50”. HISAT2 and STAR alignments of SR cDNA libraries were assembled independently with PsiCLASS v1.0.2 ^163^ with default options, and StringTie v2.2.0 ^164^ with options “-G <*ab_initio_preds*> --conservative <--rf or --fr>”. Long-read (LR) cDNA libraries were aligned with uLTRA v0.0.4 ^165^ with parameters “--ont or --isoseq, --max_intron 10000, –use_NAM_seeds” and exon hints from *ab_initio_preds*. LR cDNA alignments were cleaned and collapsed with StringTie with options “-G <*ab_initio_preds*>, -L -R”. A set of high-confidence splice junctions was selected by running Portcullis v1.2.0 ^166^ with default options on combined HISAT2 and STAR alignments. Mikado v2.3.2 ^167^ was used to generate an annotation set from only transcript evidence with options “config --mode permissive --scoring plant.yaml --copy-scoring plant.yaml --junctions <*portcullis junctions*>”, TransDecoder v5.5.0 for ORF prediction, and “serialize -- no-start-adjustment” (*rna_preds*).

Due to the scarcity of duckweed RNA-seq data from varied tissues, developmental stages, and growth conditions, we anticipated that a protein homology-based annotation approach would recover more accurate gene models for a large number of genes. The GeMoMa v1.8 ^168^ pipeline was used for this purpose. Reference proteomes from Phytozome 13 ^169^ were gathered for *A. comosus*, *A. thaliana*, *A. trichopoda*, *B. distachyon*, *N. colorata*, *O. sativa*, and *Z. marina*, and from NCBI RefSeq for *E. guineensis*. Independent GeMoMaPipeline runs were performed for each reference with default parameters except “AnnotationFinalizer.r=NO”, and then combined with GAF with the default of 10 maximum predictions per locus (*protein_preds*).

Common single-copy BUSCO v5.1.3 ^170^ sequences were determined for all novel assemblies in this study, Sp9509, and Phytozome 13 assemblies for *A. americanus*, *A. comosus*, *Z. marina* and *S. polyrhiza* 7498. MAFFT v7.487 ^171^ “--auto” was used to build protein MSAs for each BUSCO across all accessions, which were then concatenated. IQ-TREE v2.1.4 ^172^ with options “-B 1000 –mset LG,WAG,JTT” was used to build a guide tree, and a multiple whole-genome alignment was constructed with Cactus v2.0.4 ^173^. Using this alignment, a combined hints file from *ab_initio_preds*, and Lm7210 as the reference, Augustus-CGP v3.4.0 ^174^ with options “softmasking=1 --allow_hinted_splicesites=gcag,atac” was used to generate comparative gene predictions (*cgp_preds*).

To privilege the transfer of annotations from reference proteomes, while also ensuring the retention of novel gene loci predicted by other methods, the agat_sp_complement_annotations.pl script from AGAT ^175^ was run with *protein_preds* as the reference, filling in predictions at unannotated loci first with *rna_preds* and then with *cgp_preds*. MAKER v3.01.04 ^176^ was then used to evaluate the evidence supporting these complemented gene models (model_gff) both from transcript assemblies (*rna_preds*) and protein homology (*protein_preds*). A MAKER-P “standard” build was created as described in ^177^ to retain only models with evidence support or a Pfam domain. These models were further filtered to remove likely transposon sequences by screening with TEsorter v1.3.0 ^178^ against the REXdb plant TE database ^179^. If at least 25% of the amino acid sequence of any gene prediction was covered by transposase matches, and supported by fewer than 2 reference proteomes in the GeMoMa annotation, it was removed. Finally, the PASA v2.5.1 pipeline ^180^ was used as previously described ^181^ to update the gene models, using SR and LR transcript assemblies to add UTRs and alternative splice forms (*final_preds*). A subsequent round of filtering out TE and organelle-derived gene models was carried out by DC-MegaBLAST v2.13.0+ against the accession-specific organellar gene CDS and EDTA TE libraries. If more than 50% of the *final_preds* CDS sequence was covered in the top hit to either database, that gene model was removed (*final_preds_filt*).

### Phylogenetic analysis

We used OrthoFinder2 v2.5.4 ^182–184^ to infer a species tree and phylogenetic relationships among the reference and novel proteomes presented in this study (Fig 4a). A complete listing and details are provided in Table S3. *G. montanum* was used as the outgroup. The proteomes of each subgenome of the *L. japonica* hybrids were treated separately for this analysis. All vs. all alignments were computed with diamond ^185^ “--iterate --ultra-sensitive -e 0.001”. OrthoFinder was run in MSA mode “-M msa” using MAFFT v7.487 ^171^ “--localpair --maxiterate 1000” for alignments with fewer than 1,000 sequences, and default options otherwise. Trees were constructed with either IQ-TREE v2.1.4 ^172^ or VeryFastTree v3.1.0 ^186^ conditionally as follows: (species tree) “iqtree2 --alrt 1000 -T 48 - m MFP --mset Q.plant,LG,WAG,JTT”; (> 5,000 sequences) “VeryFastTree -ext AVX2 -threads 8 -double-precision”; (> 1,000 sequences) “iqtree2 -fast --alrt 1000 -T 24 -m MFP --mset Q.plant,LG,WAG,JTT”; (> 2 sequences) “iqtree2 --alrt 1000 -T 8 -m MFP --mset Q.plant,LG,WAG,JTT”. The species tree was transformed into a time tree using the make_ultrametric.py script distributed with OrthoFinder, and plotted using the phyloseq ^187^, ggtree ^188^, and deeptime ^189^ packages in R.

Beyond the individual proteomes, groupings of multiple taxa were constructed according to phylogeny (e.g. monocots, Lemnaceae, *Lemna* spp., etc.) or ecology (e.g. aquatic-floating). For each grouping, HOGs (hierarchical orthogroups) that were missing exclusively from all members of the group but not other taxa (missing HOGs), and HOGs unique to the taxa in the grouping (unique paralogs) were tabulated using R scripts. Five tables were produced in this manner reflecting different phylogenetic constraints: the “all_angiosperms” table shows missing HOGs and unique paralogs at the N1 (angiosperm MRCA) level in the species tree, with the subgenomes of the *L. japonica* hybrid accessions merged; the “ath_1mono” table shows HOGs missing from each grouping that were present in *A. thaliana* and at least one other monocot outside of the target grouping at the N4 (monocot-eudicot MRCA) level; the “ath_osa” table shows missing HOGs and unique paralogs from each grouping relative to *A. thaliana* and rice; the “intra_lemnaceae” table considers only variation within duckweeds; the “hybrids” table examines HOG variation among groupings of the *L. japonica* genomes and subgenomes. eggnog-mapper v2.1.6 ^190^ and AHRD v3.11 ^191^ were used to assign functional and Gene Ontology term annotations to the sequences in all proteomes independently, and a merged annotation record was generated for each HOG. If the HOG contained *A. thaliana*, rice, or maize sequences, symbols (from TAIR10, IGRSP-1.0, for those genes were added to the annotation. GO-term enrichment analysis, reduction, and treemap plotting were performed for each list of missing HOGs or unique paralogs under each constraint using the TopGO ^192^ and rrvgo ^193^ R packages. Significantly enriched GO terms annotated to the gene members of each HOG were added to the merged annotation (Table S3).

### Synteny analysis

Syntenic relationships between the 9 chromosome resolved Lemnaceae assemblies annotated in this study were determined and plotted using GENESPACE v1.1.4 ^194^ with “onewayBlast = TRUE”. *Z. marina* was used as an outgroup for this independent OrthoFinder run within GENESPACE, but was not used in the subsequent analysis. Lm7210 was used as the reference for riparian plots, and chromosomes of Sp9509 and Wa8730 were ordered and oriented to emphasize syntenic relationships with *Lemna* species in the plots.

### Genome size estimation by flow cytometry

Nuclear DNA content of duckweed accessions was measured by flow cytometry in triplicate, using *Spirodela polyrhiza* 7498 ^32^ and *Physalis grisea* ^195^ as controls. For each sample, two duckweed colonies from the target accession, two *S. polyrhiza* 7498 colonies and approximately 1 cm^2^ of *P. grisea* leaf were chopped with a razor blade in a 60×15mm petri dish containing 1 ml of cold Galbraith buffer (45mM MgCl_2_, 30mM sodium citrate, 20mM MOPS, 0.1% (v/v) triton X-100, pH 7.0) ^196^ for two minutes. Samples were then passed through a 30 µm CellTrics disposable filter and stained with 50 µg of Propidium iodide. Fluorescence was measured on an LSR Dual Fortessa Cell Analyzer (Becton Dickinson).

### GISH (Genomic in situ hybridization)

#### Chromosome preparation

The fronds were grown in liquid nutrient medium ^197^ under 16 h white light of 100 µmol m-2 s-1 at 24°C. The mitotic chromosome spreading was carried out according to ^198^. Fronds were treated in 2 mM 8-hydroxyquinoline, fixed in fresh 3:1 absolute ethanol: acetic acid, softened in PC enzyme mixture [1% pectinase and 1% cellulase in Na-citrate buffer, pH 4.6], macerated and squashed in 45% acetic acid. After freezing in liquid nitrogen, chromosome spreads were treated with pepsin [50 µg/ml in 0.01 N HCl], post-fixed in 4% formaldehyde in 2 × SSC [300 mM Na-citrate, 30 mM NaCl, pH 7.0], rinsed twice in 2 × SSC, 5 min each, dehydrated in an ethanol series (70, 90 and 96%, 2 min each) air-dried and inspected using spatial super-resolution structured illumination microscopy (3D-SIM) ^199^.

#### DNA isolation

For each sample, 0.3 g of fresh and healthy fronds were harvested and cleaned in distilled water, put into a 2 ml Eppendorf tube with two metal balls, frozen in liquid nitrogen, and ground by a ball mill mixer (Retsch MM400). The genomic DNA of the studied species was isolated using the DNeasy Plant Mini Kit (cat. nos. 69104-Qiagen). DNA was eluted by 200 µl buffer AE and quality checked by electrophoresis. Genomic DNA was sonicated before labeling.

#### Probe preparation

Sonicated genomic DNA (1 µg) was labeled with Cy3-dUTP (GE Healthcare Life Science) or Alexa Fluor 488-5-dUTP (Life Technologies) by nick-translation, then precipitated in ethanol ^200^ with sonicated unlabeled DNA of the other presumed parental species as carrier DNA in excess. Probe pellets from 10 µL nick translation product for GISH probes were dissolved in 100 µL hybridization buffer [50% (v/v) formamide, 20% (w/v) dextran sulfate in 2 × SSC, pH 7] at 37°C for at least 1 h. The ready-to-use probes were stored at -20°C.

#### GISH

Probes were pre-denatured at 95°C for 5 min and chilled on ice for 10 min before adding 20 µL probe per slide. Two-rounds of GISH with alternatively labeled genomic probes of the presumed parental species were performed to investigate the distribution of the corresponding probe signals on the chromosome complement of the tested clones as described ^201^.

#### Microscopy and image processing

Fluorescence microscopy for signal detection followed ^199^. To analyze the ultrastructure and spatial arrangement of signals and chromatin at a lateral resolution of ∼120 nm (super-resolution, achieved with a 488 nm laser), 3D-SIM was applied using a Plan-Apochromat 63×/1.4 oil objective of an Elyra PS.1 microscope system and the software ZENblack (Carl Zeiss GmbH).

### Data Availability

Sequencing datasets and genome assemblies generated during the current study are available at NCBI (GEO SuperSeries: GSE238136, BioProject: PRJNA999459). Genome assemblies and annotations presented here are available along with browsing and analysis tools at www.lemna.org. Code is available at https://github.com/martienssenlab/lemnaceae-genomes-manuscript. All materials are available upon request.

## Acknowledgements

We thank our colleagues in the duckweed community for their many contributions and enthusiastic support. This work was primarily supported by Howard Hughes Medical Institute (R.A.M.) and by the U.S. Department of Energy, Office of Science, Office of Biological and Environmental Research program under Award Number DE-SC0018244 (E.E., K.A., B.P., C.M-E, U.R., E.L., R.A.M.) as well as the Foundation for Food and Agricultural Research, Seeding Solutions Grant CA21-SS-0000000100 (E.E., C.M-E., U.R., R.A.M). In addition, this work was supported by a Hatch project (12116) and a Multi-State Capacity project (NJ12710) from the New Jersey Agricultural Experiment Station at Rutgers University (K.A., B.P., E.L.) and by the Tang Genomics Fund (N.H., B.A., K.C., A.A., T.P.M.). This work was performed with assistance from the US National Institutes of Health Grant NIH S10OD028632-01, which supports the HPC cluster at Cold Spring Harbor Laboratory.

## Author contributions

Conceptualization, E.E., T.P.M., I.S., E.L., and R.A.M.; Investigation, E.E., B.A., K.A., P.T.N.H., C.M-E., V.S., B.P., K.C., A.A., S.C.L., U.R., J.A.B.; Formal Analysis, E.E., B.A., K.A., V.S., K.C., A.A., I.S., T.P.M., and R.A.M.; Data Curation, E.E.; Writing - Original Draft, E.E., T.P.M., and R.A.M.; Writing - Review & Editing, E.E., J.A.B., I.S., E.L., T.P.M., and R.A.M.; Visualization, E.E., C.M-E., V.S., T.P.M., and R.A.M.; Project Administration, E.E. and R.A.M.; Funding Acquisition, E.E., E.L., T.P.M., and R.A.M.

