## Supplemental Figures for "The genomes and epigenomes of aquatic plants (Lemnaceae) promote triploid hybridization and clonal reproduction"

### Title

### Author affiliations and footnotes

[1]*Howard Hughes Medical Institute, Cold Spring Harbor Laboratory, Cold Spring Harbor, NY, USA*
[2]*The Plant Molecular and Cellular Biology Laboratory, The Salk Institute for Biological Studies, La Jolla, CA 92037, USA*
[3]*Department of Plant Biology, Rutgers, The State University of New Jersey, New Brunswick, NJ, USA*
[4]*Leibniz Institute of Plant Genetics and Crop Plant Research (IPK), Gatersleben, D-06466 Stadt Seeland, Germany*
[5]*Biology Faculty, Dalat University, District 8, Dalat City, Lamdong Province, Vietnam*
[6]*Department of Marine Sciences, Faculty of Fisheries and Marine Sciences, Universitas Padjadjaran, Bandung 40600, Indonesia*
[7]*Biological Sciences, University of Missouri at Columbia, Columbia, MO, USA*
[8]Lead contact
[*]Corresponding author

### Contact information

Robert A. Martienssen:
Todd P. Michael:
Eric Lam:

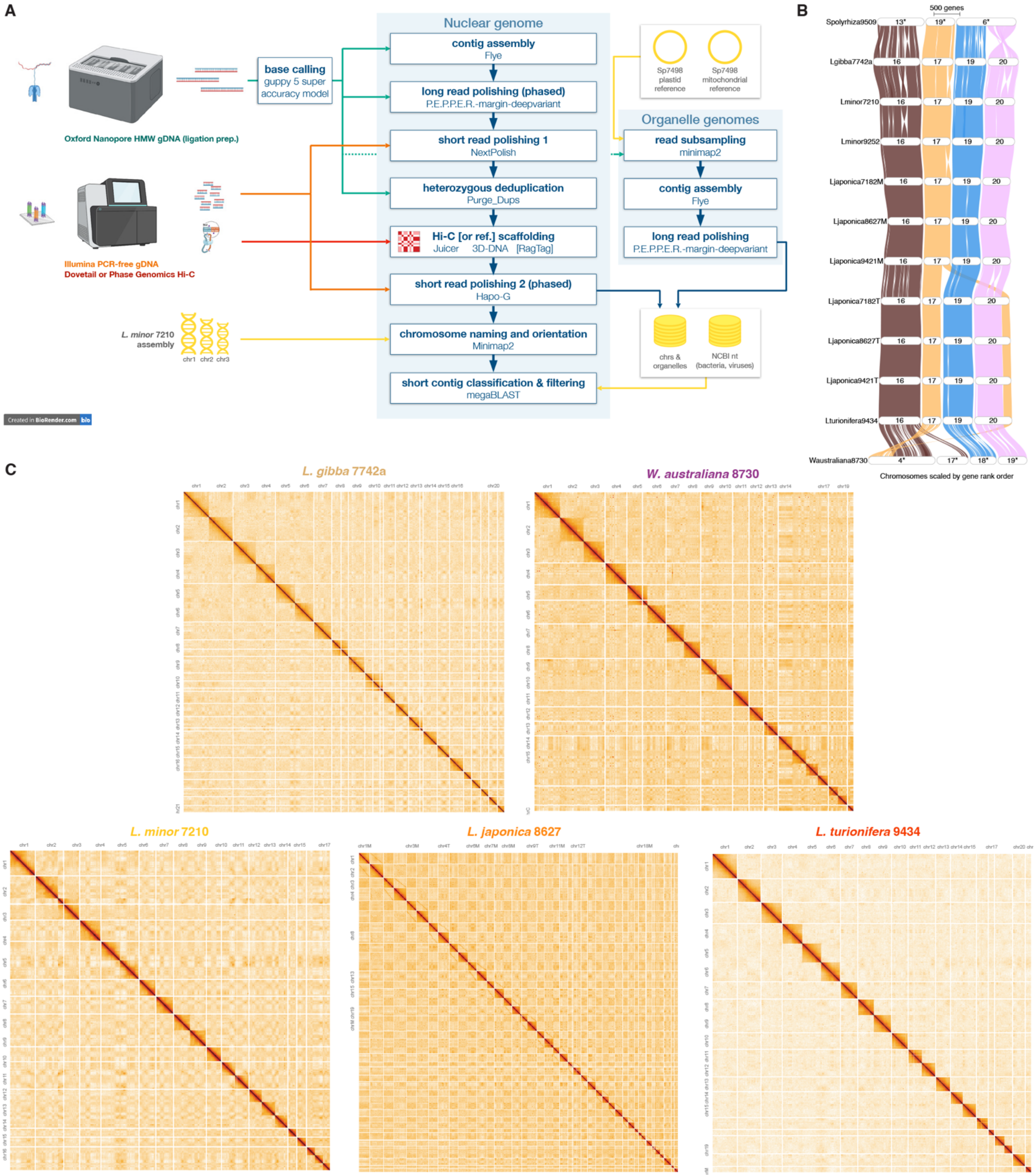


**Figure S1. Assembly strategy and structural details of Lemnaceae genomes, related to Figure 1. (A)** Genome assembly pipeline. Input data types are shown with color coding corresponding to the arrows that indicate their use throughout the pipeline. **(B)** Detail of the large-scale translocation detected in all *L. turionifera* genomes involving the arms of *Lemna* Chromosomes 17 and 20. Ribbons show blocks of syntenic genes. Twists indicate relative inversions. **c** Genome-wide Hi-C chromatin contact maps. Contact probabilities are shown after balancing with the ICE method and vizualization with HiGlass at 1 Mbp resolution with a logarithmic color scale. Chromosomes are ordered from left to right (Chr1M, Chr1T...Chr21M, Chr21T for *L. japonica*).

**A**

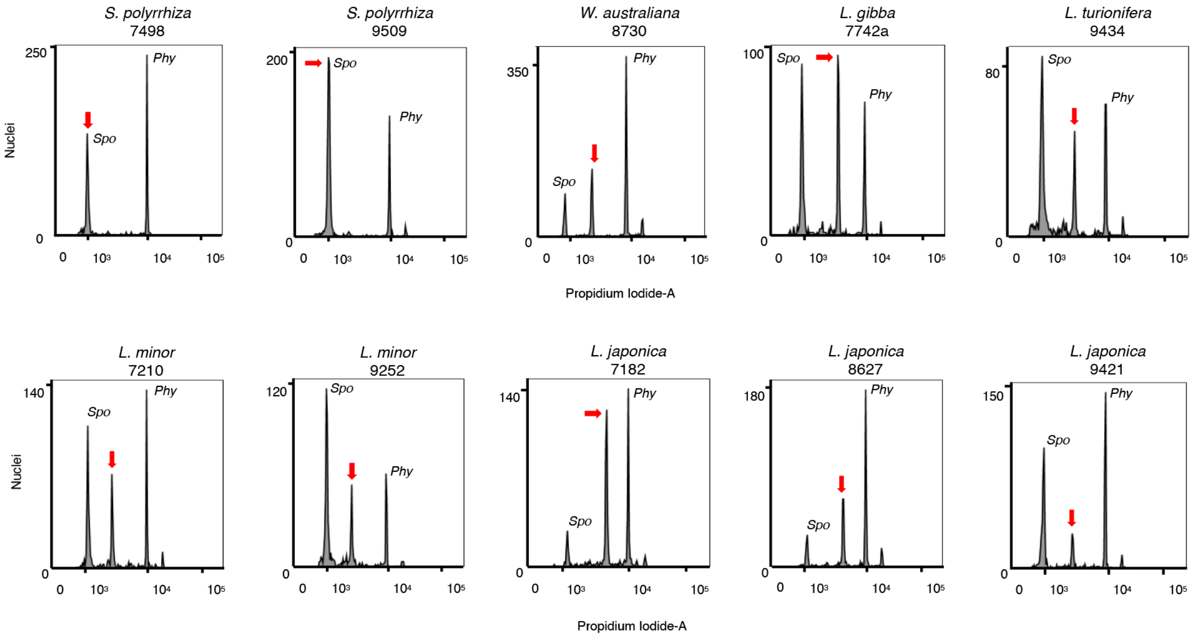


**B**

| | *S. polyrrhiza* 7498 (2n = 2x) | *P. grisea* (2n = 2x) | *S. polyrrhiza* 9509 (2n = 2x) | *L. gibba* 7742a (2n = 2x) | *L. minor* 7210 (2n = 2x) | *L. minor* 9252 (2n = 2x) | *L. japonica* 7182 (2n = 3x = MTT) | *L. japonica* 8627 (2n = 3x = MMT) | *L. japonica* 9421 (2n = 2x = MT) | *L. turionifera* 9434 (2n = 2x) | *W. australiana* 8730 (2n = 2x) |
|---|---|---|---|---|---|---|---|---|---|---|---|
| **2C DNA content (Mbp)** | 301 ± 4 | 2876 ± 41 | 316 ± 0 | 1041 ± 4 | 758 ± 13 | 762 ± 2 | 1244 ± 8 | 1157 ± 13 | 860 ± 11 | 901 ± 13 | 772 ± 2 |

**Figure S2. Genome size measurements, related to Figure 2. (A)** Flow cytometry fluorescence measurements of nuclei stained with propidium iodide. Red arrows indicate the peak corresponding to the Lemnaceae analyte in each sample. Spo = low standard, *Spirodela polyrhiza* 7498, 2C = 316; Phy = high standard, *Physalis grisea*, 2C = 2,740. **(B)** Estimates of genome size calculated from **(A)** using both standards. Reciprocal estimates of the 2C DNA content of the standards (*) as measured in this study are shown for reference.

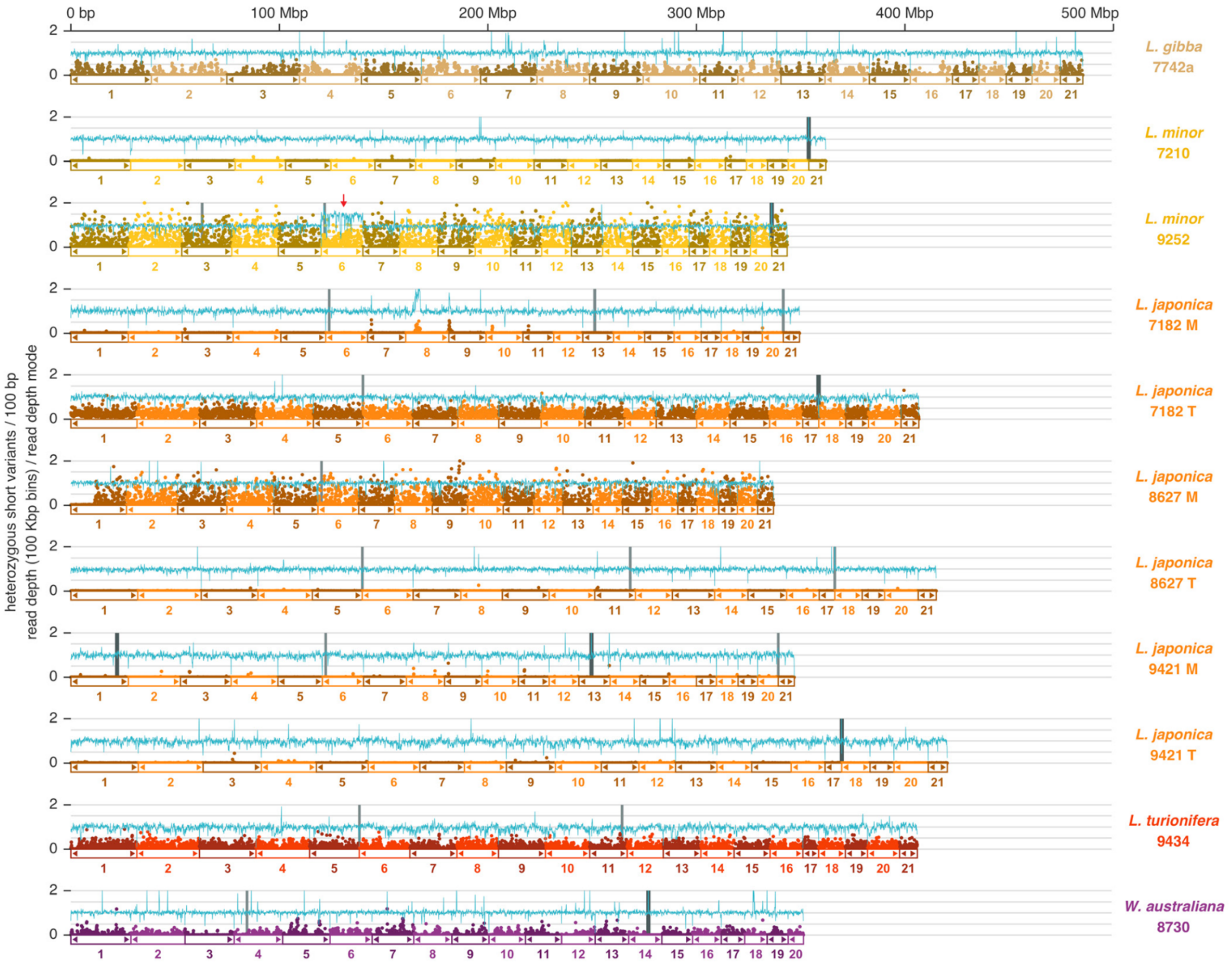


**Figure S3. Variations in heterozygosity, rDNA location, and chromosome dosage, related to Figure 1.** Genomic read coverage and variants across assembled pseudomolecules. Long read coverage (mapQ >= 30) normalized to the mode of read depth across each subgenome assembly is shown as a light blue horizontal line. Counts of heterozygous short variants in 100bp bins (SNVs + INDELs < 50 bp) are shown as colored dots. 25S rDNA loci are shown as blue vertical bars, with the most conserved loci in a darker shade. Arrows beneath are shown where telomeric repeat sequence was present in the assembly within 10 Kbp from the end of the scaffold. Trisomy of Lm9252 Chromosome 6 is indicated by a red arrow.

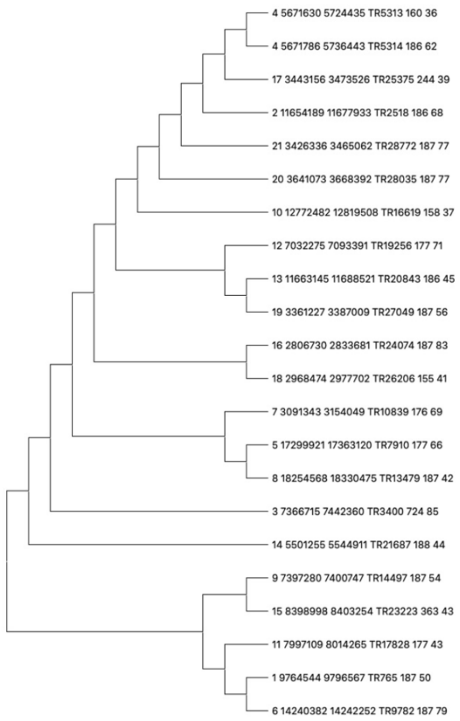


**Figure S4. Phylogenetic analysis of centromere tandem arrays, related to Figure 3.** Phylogenetic analysis of the *L. minor* centromere tandem array. The 154, 177, and 187 bp tandem arrays were extracted from the *L. minor* genome and aligned to one another to see if they were distinct arrays and if they were found on separate chromosomes. Name: chromosome number, start position, stop position, tandem array name, tandem array length, and identity to the predicted tandem array.

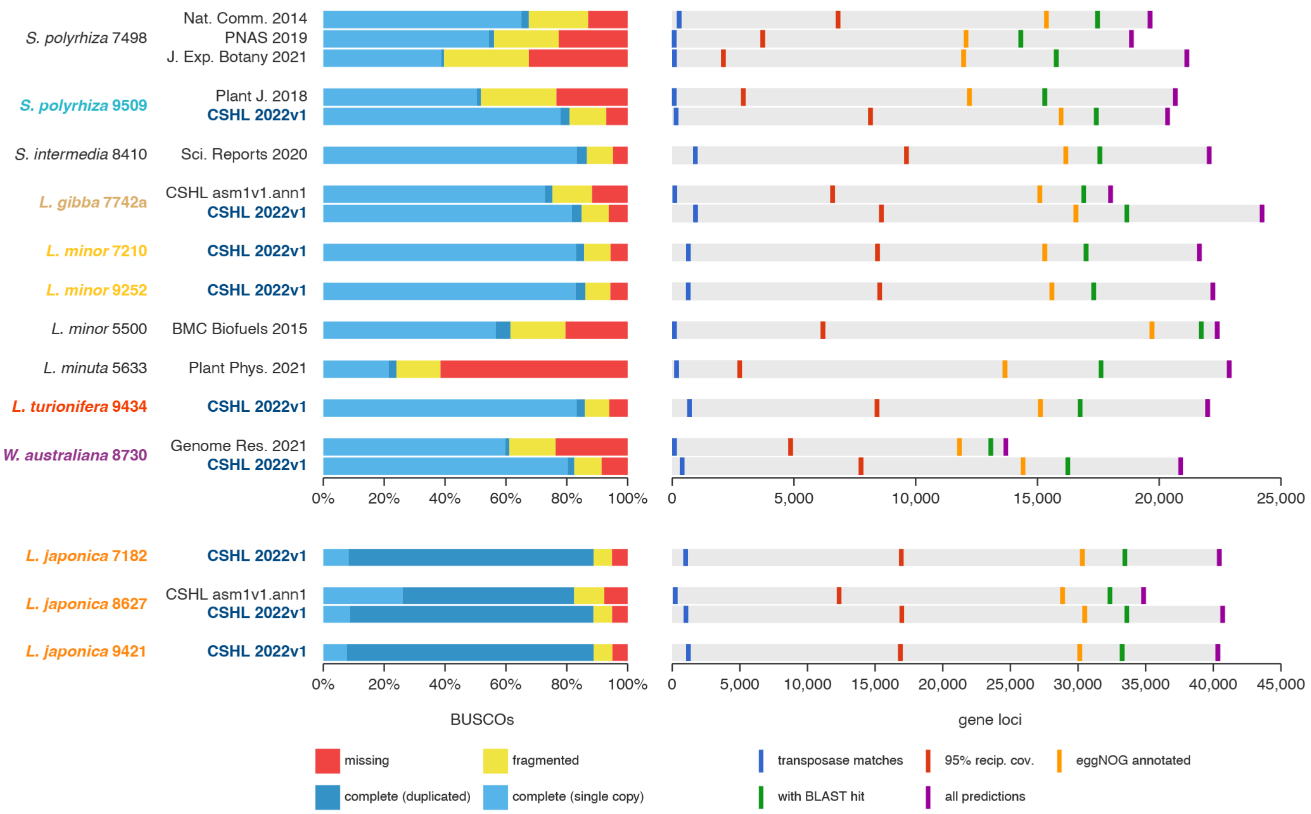


**Figure S5. Protein coding gene annotation and quality assessment, related to Figure 4.** Protein coding gene annotation quality assessment by BUSCO v5 (left) and BLAST alignment (right). For some accessions, previously published annotations are compared with those from this study (CSHL 2022v1). BUSCO protein mode analysis was used with the liliopsida ODB10 database (n=3236). For each proteome annotation, sequences were: (1) screened for transposase sequence similarity and domains using TEsorter; (2) aligned with Diamond BLASTP to a database of the following proteomes: UniProt/SwissProt Plants, *Acorus americanus*, *Arabidopsis thaliana*, *Amborella trichopoda*, *Musa acuminata*, *Nymphaea colorata*, *Oryza sativa*, *Zostera marina*, and *Elaeis guineensis*; (3) realigned with hits requiring 95% coverage of both query and target; (4) annotated with EggNOG mapper against the EggNOG v5 database.

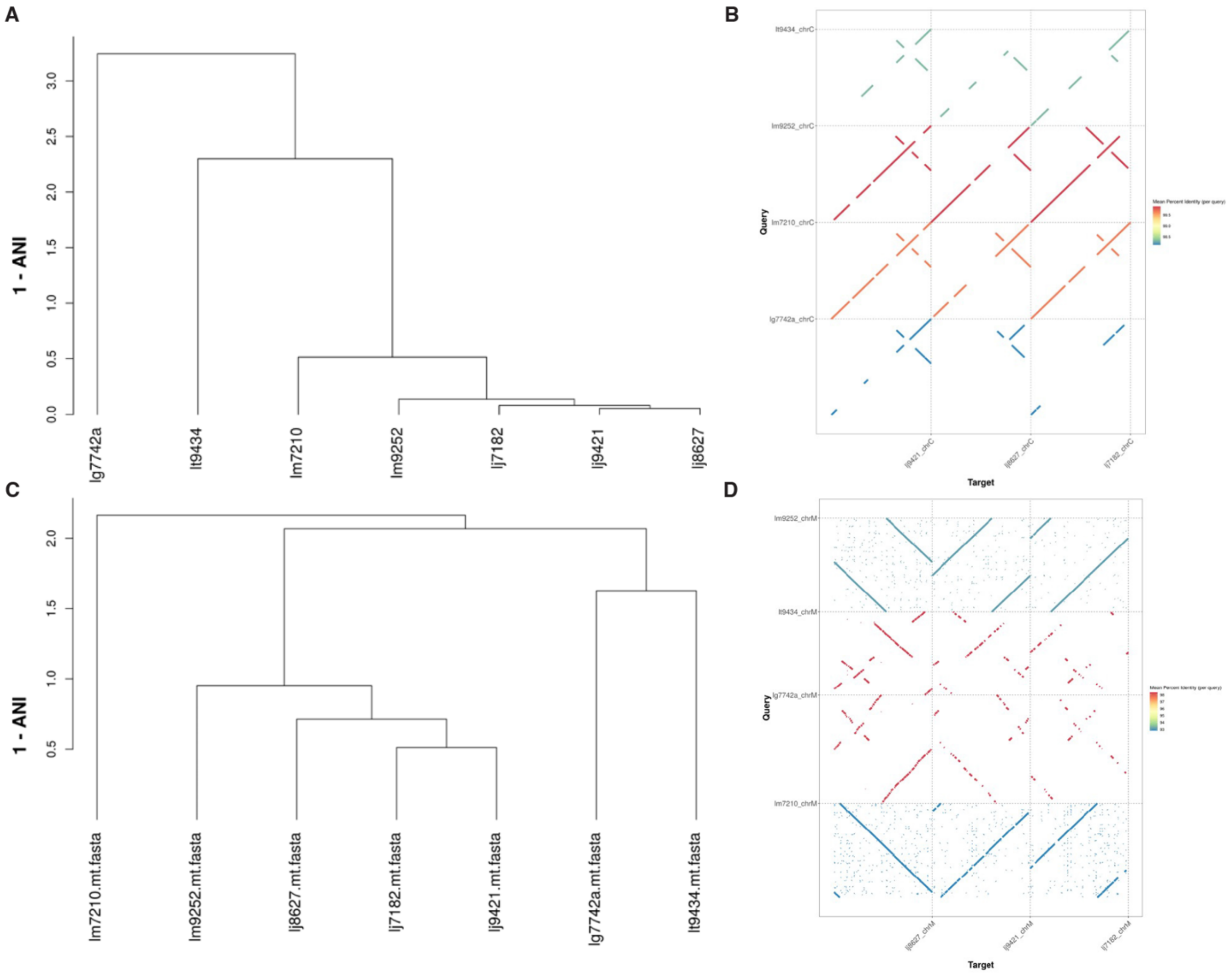


**Figure S6. Phylogenetic comparison of Lemna organelle genomes, related to Figure 2. (A,C)** Phylogenetic trees of Lemnaceae organelle genomes in this study drawn from the the average nucleotide identity (ANI) whole genome similarity metric. **(B,D)** Dotplots of organelle genome pairwise alignments. **(A,B)** Plastid genomes. **(C,D)** Mitochondria genomes.

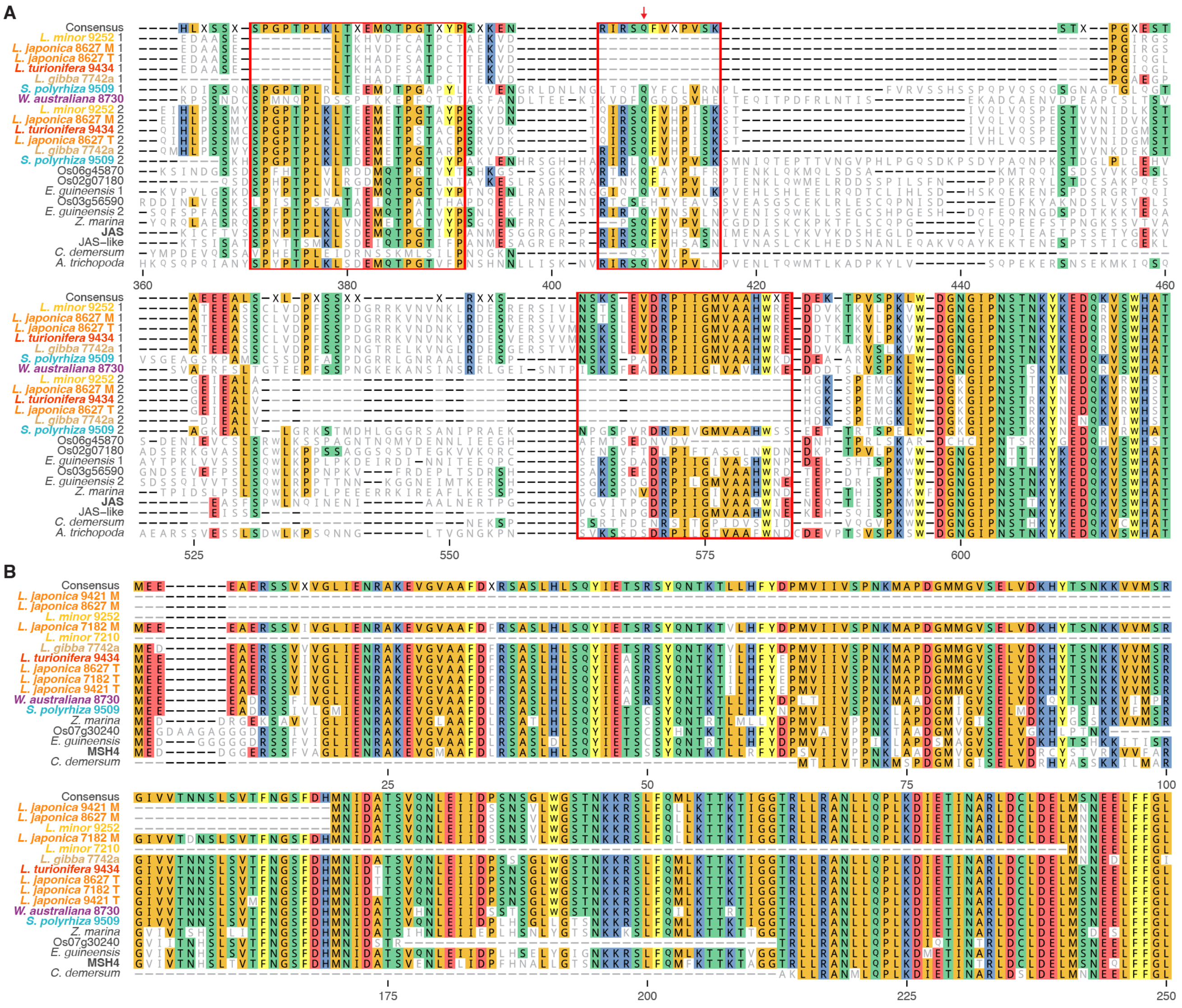

**Figure S7. Mutations in meiotic genes, related to Figures 4 and 6. (A)** Selected regions of a MAFFT amino acid multiple sequence alignment of JAS (At1g06660) and JAS-like (At2g30820) orthologs identified in a subset of Lemnaceae and other accessions. Multi-residue deletions and substitutions in well-conserved regions are outlined in red. Gene locus identifiers are shown only for rice and *Arabidopsis* for brevity, and multiple orthologs or paralogs belonging to a single accession are denoted by an arbitrary numeric suffix. Location of the *Arabidopsis jas-1* EMS mutant allele inducing an early stop codon is shown with a red arrow. **(B)** Selected regions of a MAFFT alignment of *MSH4* (At4g17380) ortholog amino acid sequences, showing N-terminal truncations in most *Lemna minor* orthologs.
