## Supplementary material for "The genomes and epigenomes of aquatic plants (Lemnaceae) promote triploid hybridization and clonal reproduction": Table S1

### Table S1. Assembly and annotation statistics of chromosome-resolved Lemnaceae genomes, related to Figure 1.

| Genomes | *S. polyrhiza** greater duckweed **Sp9509** v3 JCVI | *W. australiana* Australian watermeal **Wa8730** v3 CSHL | *L. gibba* swollen duckweed **Lg7742a** v3 CSHL | *L. minor* lesser duckweed **Lm7210** v1 CSHL | *L. minor* lesser duckweed **Lm9252** v1 CSHL | *L. japonica* 2n *L. min.* x 1n *L. tur.* **Lj8627** v3 CSHL | *L. japonica* 1n *L. min.* x 2n *L. tur.* **Lj7182** v1 CSHL | *L. japonica* 1n *L. min.* x 1n *L. tur.* **Lj9421** v1 CSHL | *L. turionifera* turion duckweed **Lt9434** v1 CSHL |
|---|---|---|---|---|---|---|---|---|---|
| chromosome # | 2n = 40 | 2n = 40 | 2n = 42 | 2n = 42 | 2n = 42 | 3n = 63 | 3n = 63 | 2n = 42 | 2n = 42 |
| **Scaffolds** | | | | | | | | | |
| span (Mbp) | 139 | 355 | 491 | 363 | 354 | 772 | 764 | 771 | 410 |
| # > 1Mbp | 20 | 20 | 21 | 21 | 21 | 42 | 42 | 42 | 21 |
| longest (Mbp) | 12 | 29 | 39 | 29 | 28 | 32 | 32 | 32 | 32 |
| gap bases (%) | 0.02 | 0.01 | 0.03 | 0.00 | 0.01 | 0.04 | 0.00 | 0.00 | 0.02 |
| **Contigs** | | | | | | | | | |
| # | 97 | 229 | 474 | 85 | 554 | 1,148 | 561 | 373 | 269 |
| N50 (Mbp) | 2.9 | 5.2 | 3.2 | 13.9 | 7.0 | 3.2 | 6.2 | 5.6 | 5.3 |
| **Repeats** | | | | | | | | | |
| all interspersed (%) | 12 | 48 | 71 | 59 | 57 | 62 | 62 | 63 | 61 |
| intact LTR-RT/Mpb (#) | 0.1 | 0.6 | 2.0 | 1.7 | 1.2 | 1.3 | 1.3 | 1.6 | 1.3 |
| **Genes** | | | | | | | | | |
| Protein coding | 20,342 | 20,882 | 24,222 | 21,649 | 22,205 | 40,692 | 40,444 | 40,336 | 21,988 |
| transcribed ncRNA | 647 | 865 | 1,852 | 888 | 1,252 | 2,100 | 2,095 | 2,107 | 976 |
| snoRNA/snRNA | 541 | 864 | 1,141 | 785 | 751 | 2,017 | 2,016 | 2,286 | 1,210 |
| tRNA | 237 | 623 | 470 | 260 | 253 | 486 | 482 | 479 | 278 |
| miRNA loci | 59 | 65 | 74 | - | - | 108 | - | - | - |

* = prior assembly reannotated in this study.
